# EpiPred: A gene-specific machine learning model for classifying missense variants in the epilepsy-related gene *STXBP1*

**DOI:** 10.1101/2025.08.09.669488

**Authors:** Jeffrey D Calhoun, Chengbing Wang, Carina Biar, Jonathan R Gunti, John Lee, Aaron M Geller, Jung H Hong, Santiago Schnell, Louis T Dang, Yu Wang, Jack Parent, Lori L. Isom, Michael D Uhler, Heather C. Mefford, M. Elizabeth Ross, Vanessa Aguiar-Pulido, Gemma L Carvill

**Author notes:** Correspondence can be addressed to Gemma L. Carvill and M. Elizabeth Ross. Co-senior authors (M. Elizabeth Ross, Vanessa Aguiar-Pulido, Gemma L Carvill). These authors contributed equally to this work (Jeffrey D Calhoun, Chengbing Wang).

## Abstract

Missense variants in the STXBP1 gene are a frequent cause of early-onset developmental and epileptic encephalopathies and related neurodevelopmental disorders, but the clinical interpretation of these variants remains a major challenge. Most reported STXBP1 missense variants are classified as variants of uncertain significance (VUS), complicating diagnosis, counseling, and patient eligibility for precision therapies. Here, we developed EpiPred, a gene-specific machine learning classifier that predicts the pathogenicity of STXBP1 missense variants by integrating computational features with empirical evidence from cellular assays. Trained on a curated set of pathogenic and benign variants, EpiPred outperformed leading global prediction tools in accuracy, sensitivity, and specificity. We validated the model’s predictions using functional assays that measure protein abundance, solubility, stability, and interaction with the SNARE complex partner syntaxin 1. These biochemical readouts aligned closely with model outputs and enabled reclassification of several likely misdiagnosed variants. We deployed EpiPred in an interactive web application that allows clinicians, researchers, and patients to explore predictions for all possible missense variants in STXBP1. Our approach illustrates the power of gene-specific predictive modeling combined with experimental validation to improve variant interpretation and diagnostic resolution. By identifying likely pathogenic STXBP1 variants, including those that may respond to emerging therapies such as molecular chaperones, EpiPred supports more precise genetic diagnoses and offers a generalizable framework for other clinically relevant genes in neurological disease.

**ONE SENTENCE SUMMARY:** EpiPred improves STXBP1 variant interpretation, enabling precision genetic diagnoses and promoting access to targeted precision therapies

## INTRODUCTION

The rare, pediatric-onset genetic epilepsies are characterized by recurrent, medication refractory seizures, comorbid developmental delay, and behavioral challenges that often occur in the context of the developmental and epileptic encephalopathies (DEEs). The prevailing genetic etiology of these epilepsies is now well established, with more than half of individuals receiving a genetic diagnosis via gene panel or exome sequencing(*1–4*). However, a major consequence of widespread genetic testing is the sharp increase in variants of uncertain significance (VUS), particularly missense variants whose effects are challenging to interpret. VUS lack sufficient information to interpret as pathogenic or benign and present a significant challenge for families and clinicians alike. There are currently over 1.1 million VUS in ClinVar, the database for deposition of clinically-relevant genetic variants, accounting for over 90% of all variants in this collation(*5*). Moreover, roughly half of all clinically reported variants are VUS(*6*).

Among epilepsy-related genes, STXBP1 stands out as a high-priority candidate for gene-specific pathogenicity modeling. It is one of the most commonly implicated genes in DEEs, yet displays diffuse missense variation across its coding sequence, lacks well-defined functional domains, and harbors a disproportionately high number of missense VUS. Moreover, there is a growing body of functional data, ranging from abundance and solubility to protein-protein interactions, that provides a robust empirical foundation for variant classification. These properties make STXBP1 both clinically significant and mechanistically tractable for the development of a tailored, gene-specific variant effect predictor. Here, we leverage this opportunity to build EpiPred, a supervised machine learning model designed to classify STXBP1 missense variants using both computational and biochemical characterization. This study forms part of the Epilepsy Multiplatform Variant Prediction (EpiMVP) study, an NIH-funded, multicenter collaborative effort to resolve epilepsy-related VUS through the integration of computational modeling and cellular and animal-based functional assays.

There is a strong clinical need for STXBP1 missense VUS resolution. Clinically, disease-causing *STXBP1* variants were first described in the DEEs(*7*), but there has been a widening of the phenotypic spectrum to neurodevelopmental disorders (NDDs) more broadly. However, epilepsy remains a prominent feature, present in 95% of patients(*8*). Genetic interrogations in ∼13,000 individuals with epilepsy revealed *STXBP1* as the fifth most common epilepsy-related gene; only pathogenic variants in the ion channels (*SCN1A/2A*, *KCNQ2*) and *CDKL5* were more common(*1*). Moreover, a meta-analysis of all *de novo* variants linked to human disease showed *STXBP1* to be the sixth most commonly mutated gene relative to its size(*9*). *STXBP1* has been included in gene panels since ∼ 2011, and as such there are hundreds of VUS in the gene. For example, in a cohort of ∼10,000 individuals sequenced for *STXBP1,* an almost equal number of pathogenic variants (0.48%) were identified as the number of individuals reported with an *STXBP1* VUS (0.3%)(*4*). When we began this study there were 150 missense VUS listed in ClinVar; this number has subsequently risen to 222 an increase of 33% in just a few years, highlighting the need for VUS resolution.

The majority of pathogenic *STXBP1* variants are truncations which arise *de novo*, consistent with a disease mechanism of haploinsufficiency. STXBP1 lacks any distinct functional domains, and thus perhaps unsurprisingly, the over 100 pathogenic missense variants are located throughout the protein. This lack of aggregation into discrete functional domains limits VUS resolution strategies that have been shown to be effective in other epilepsy-related genes, such as the voltage-gated ion channels, where variants tend to aggregate in the pore-forming regions(*10*). Thus, we must leverage other key features of pathogenic missense variants to discriminate between pathogenic and benign *STXBP1* variants. STXBP1, also known as UNC18/MUNC18, together with STX1, synaptobrevin and SNAP25 form the SNARE complex(*11*). This complex is a key component of neuronal function, as it facilitates fusion of the synaptic vesicle with the plasma membrane allowing neurotransmitter release into the synapse. Fortunately, there is an increasing understanding of how missense pathogenic variants disrupt STXBP1 function and cause disease. This includes the reduced abundance of mutant STXBP1 proteins in HEK293 cells, *Stxbp1* null mouse neurons(*12, 13*), induced pluripotent stem cells (iPSCs)(*14*), as well as engineered *c. elegans* models (*15–17*). Pathogenic missense variants also lead to reduced interaction with SNARE partner, STX1, and interactors Doc1(*12, 13*). Finally, there is some evidence for protein aggregation (*13, 17*). Together, these studies converge on a common STXBP1 missense phenotype characterized by reduced protein abundance, stability, and solubility, along with disrupted STX1 interaction—features that enable the functional resolution of missense VUS.

Robust functional assays enable rigorous VUS resolution; however, few studies have used multiple benign missense variants to truly benchmark assays. Here we leverage these functional readouts to test the accuracy of global and gene-specific variant effect predictors (VEPs) to predict the effects of *STXBP1* missense variants using robust positive (pathogenic) and negative (benign) controls. Global, or genome-wide, VEPs, such as CADD, MCAP, REVEL and alphaMissense, ClinPred, VARITY and BayesDel are often used as supporting evidence when determining variant pathogenicity as per the American College of Medical Genetics (ACMG) guidelines(*18, 19*)(*20–22*). These global predictors are the most common and generalize reasonably well due to the large datasets available for training. However, there are increasing efforts to develop both disease and gene-specific classifiers, which generalize poorly, but can offer improved performance in a clinical context. Examples include cancer (Cancer-specific High-throughput Annotation of Somatic Mutations (CHASM)) and cardiac (CardioBoost) condition specific VEPs, which offer increased positive predictive values and 4-24% improvement over global-VEPs (*23, 24*). There are also a few limited examples of gene-specific classifiers, mostly in cancer. In particular, for the major breast and ovarian cancer driver gene(s), *BRCA1/2*, there are four gene-specific VEPs that outperform global VEPs(*25–28*), demonstrating the potential utility of this approach.

Here, we generated an *STXBP1*-specific VEP, EpiPred (Epilepsy Prediction), using a supervised machine learning approach. We benchmarked EpiPred-*STXBP1* against commonly used global VEPs and tested our predictions using known readouts of *STXBP1* pathogenicity (**Figure 1A**). Collectively, EpiPred-STXBP1 exhibited improved performance over global VEPs and had high concordance with experimental analyses across 20 variants. We also generated predictions for every possible missense variant in *STXBP1* and demonstrate that the core of the STXBP1 protein is susceptible to perturbation by missense variants and likely underpins reduced abundance of STXBP1. Importantly, the reduced abundance of STXBP1 can be rescued in both animal and cellular models using 4-phenylbutyrate (PBA), an FDA-approved medication commonly used to lower serum ammonia levels in individuals with urea cycle disorders (*17*). A small clinical trial of PBA in individuals with *STXBP1*-related neurodevelopmental disorders suggests that the treatment may reduce seizure frequency and improve EEG abnormalities (*29*). Thus, collectively our study highlights the power of integrating computational and functional characterization of STXBP1 missense variants to inform precision diagnostics and empower precision clinical care.

**Figure 1:**
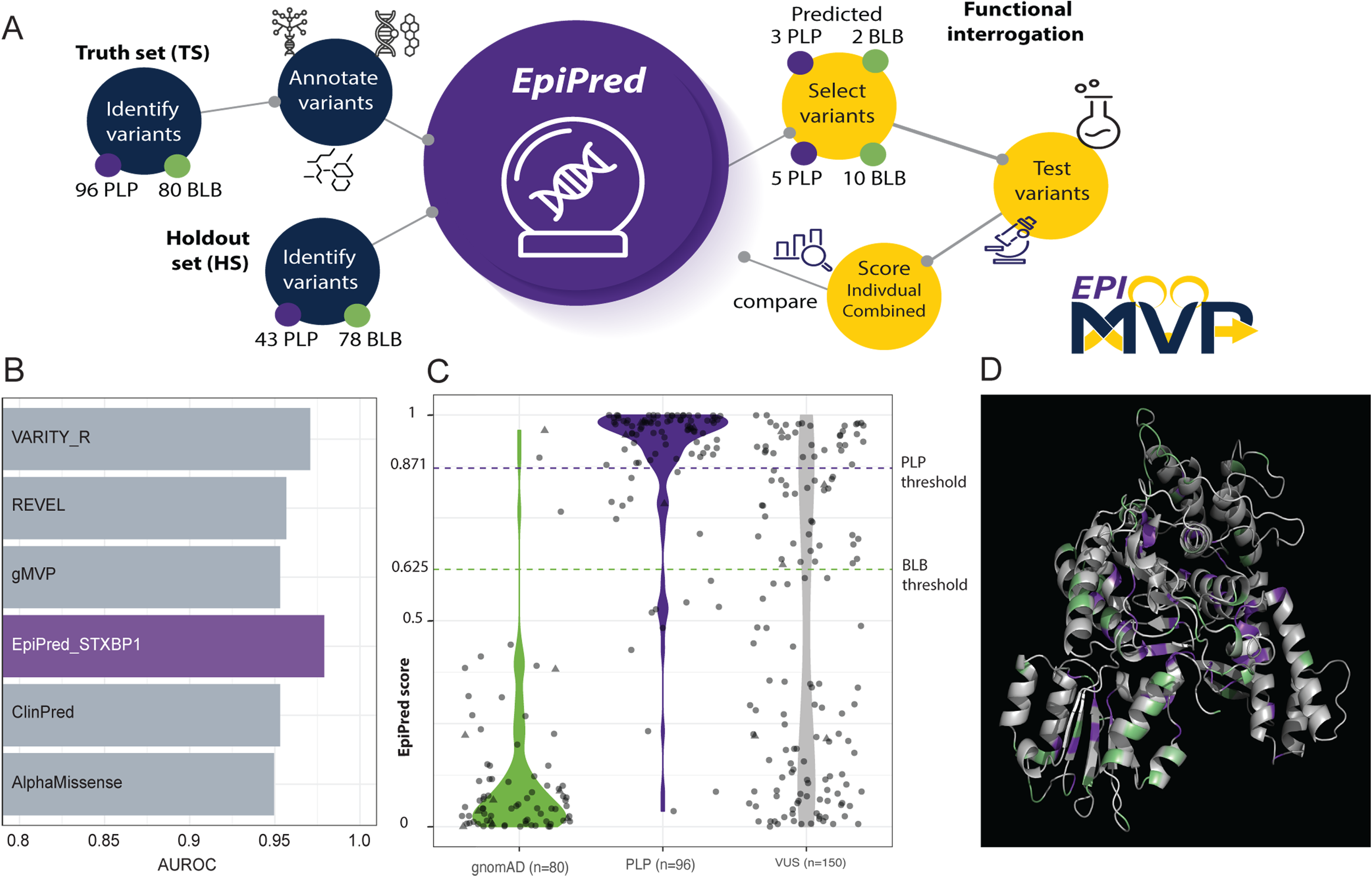
Supervised learning facilitates the generation of an STXBP1-specific variant effect predictor, EpiPred. (**A**) Overview of the approach adopted for the generation of the EpiPred model, including the input datasets (blue) and the functional testing of the EpiPred predictions using three biochemical assays (stability, abundance, solubility) and STX1 interaction in an overexpression model system (yellow). (**B**) Comparison of area under the receiver operator curve (AUROC) for EpiPred and a subset of high-performing globally-trained VEPs across the full truth set. (**C**) EpiPred Probability of PLP (EpiPred-P^PLP^) score distributions in the full truth set separated by class as well as the full set of VUS. EpiPred-P^PLP^ scores of 1 suggest pathogenicity, while scores of 0 suggest neutrality (**D**) EpiPred-P^PLP^ output for known PLP and BLB variants used in the truth set mapped onto hSTXBP1 predicted structure (AlphaFold structure: AF-P61764-F1-model_v4), EpiPred-P^PLP^ scores close to 1 are shown in purple, while scores close to zero appear in green.

## METHODS

### Supervised learning to develop EpiPred-STXBP1

To develop our training set, missense variants were curated from multiple sources including ClinVar, gnomAD v3 (variants with an allele count >3), the scientific literature, and in-house databases from clinical genetic testing companies (Geisenger, GeneDx, Invitae). After removing duplicates, our dataset contained 80 benign or likely benign (BLB) variants (gnomAD and ClinVar) and 96 pathogenic or likely pathogenic (PLP) variants (ClinVar) defined as per ACMG criteria. This training dataset is referred to as the Truth Set (TS) (**Table S1**). We further identified 150 VUS, listed as either VUS or conflicting in ClinVar (**Table S2**). Each variant was annotated with 42 features including amino acid properties, conservation, (n=42; **Table S3**) using either Variant Effect Predictor or custom scripts. Empty values were replaced with the median value for that feature. Initially, multiple models including multiple linear regression, decision tree, and random forest were explored. We identified random forest as a model which could accurately perform the classification task, in agreement with previous gene-specific VEPs(*10*). A random forest model was developed using the Python scikit-learn package and hyperparameters were tuned using five-fold cross validation. EpiPred-*STXBP1* was trained using 160 trees with maximum depth 5 on a random 60% subset of the TS and tested on the remaining 40%. Performance of EpiPred-*STXBP1* and competing *in silico* pathogenicity predictors were investigated using the R pROC package.

### Validation of EpiPred-STXBP1 on a holdout dataset

Missense variants were curated from gnomAD v4, the Regeneron million exome server, and the literature.(*30*) For BLB variants, we required an allele count of >2 in either gnomAD v4 or the Regeneron exome server. Variants that overlapped with the TS were removed. The final Holdout Set (HS) contained 121 unique missense variants, including 78 BLB and 43 PLP (**Table S8**).

### Gaussian Mixture Modeling

The mclust R package was used to develop a univariate, unequal variance model based on the EpiPred scores for TS and HS variants(*31*). We found the upper 95% mean confidence interval for BLB variants was 0.625179 while the lower 95% mean confidence interval was 0.8711709. Based on this modeling, we considered variants with EpiPred scores in the interval between these two values to be ambiguous.

### Global and local model explanations

To investigate global and local model decisions, we utilized a tree explainer within the Python SHapley Additive exPlanations (SHAP) package(*32*). At the global level we used these interpretations to determine what features drove EpiPred generally across the STXBP1 missense variant landscape. We leveraged local explanations, i.e. for each variant, in particular those with conflicting EpiPred predictions to determine whether certain features drove these misclassifications.

### Functional characterization of STXBP1 missense variants

A subset of variants (n=20) was selected for functional validation; experimenters were blinded to the classification (PLP/VUS/BLB) of each variant. At least three independent replicates were tested for each variant. Moreover, all quantification was performed by two independent experimenters blinded to variant identity. t-tests were performed for all pairwise comparisons of each variant to wildtype, significance threshold was 0.05.

The plasmid (pPB-CAG-EGFP Addgene 40973), with the EGFP coding region replaced with either wildtype *STXBP1* or one of the missense variants, was transformed into DH5α Competent Cells (18265017, Thermo Fisher Scientific). Three bacterial clones were picked and inoculated into LB broth with 100µg/ml carbenicillin. The bacteria were cultured for 16-18 hrs. The plasmid DNA was prepared with GenCatch™ Plasmid DNA Mini-Prep Kit (2160-250, Epoch Life Science, Inc).

#### STXBP1 abundance assay

HEK293T cells were maintained with Corning® DMEM [+] 4.5 g/L glucose, L-glutamine, sodium pyruvate (45000-304, VWR), supplemented with 10% Fetal Bovine Serum, value (A5256701, Thermo Fisher Scientific) and 100 U/ml Penicillin-Streptomycin (10,000 U/mL) (15140122, Thermo Fisher Scientific). The day before transfection, 0.5×10^6^ cells/well were plated into 6-well plates. The cells were transfected with 1 ug of the same batch-prepared plasmid with PolyJet™ In Vitro DNA Transfection Reagent (SL100688, SignaGen Laboratories). 48-hr post-transfection, cells were washed with cold 1XPBS and lysed using RIPA buffer containing protease inhibitors. The protein concentration was determined with Pierce™ BCA Protein Assay Kits (23227, Thermo Fisher Scientific). Equal amounts of protein were separated by SDS-PAGE and transferred to polyvinylidene difluoride (PVDF) membranes (1620177, Bio-Rad Laboratories, Inc.). After blocking, blots were probed with Munc18-1 (D406V) Rabbit mAb #13414 (1:1000, Cell Signaling Technology). Blots were stripped and reprobed for Alpha Tubulin with Alpha Tubulin Monoclonal antibody (1:2000, 66031-1-Ig, Proteintech Group).

#### STXBP1 solubility assay

HEK293T cells in 60mm Petri dishes were transfected with different amounts of plasmid DNA to ensure a similar protein abundance, according to the plasmid DNA expression potency. 48 hrs post-transfection, the cells were washed with PBS + 1mM MgCl2. After that, 1XPBS+0.1% Triton X 100+ protease inhibitors+ Pierce™ Universal Nuclease for Cell Lysis (1:1000, 88700, Thermo Scientific) was added to the dishes. The cells were collected into 1.6ml Eppendorf tubes with a cell scraper. The tubes were placed on an agitation shaker and shaken gently in the cold room for 1hr. The lysates were pelleted at 4°C, 20 000 g for 20min. The supernatants (containing the soluble STXBP1 protein) were transferred into new Eppendorf tubes and placed in ice. The pellets were resuspended with 1% SDS in PBS. The tubes were placed on a shaker at 30°C at 300 rpm, for 20 min, and then centrifuged at 10 000 g for 20min at 23°C. The pellets were removed. The supernatants stood at room temperature during the protein concentration measurement with a BCA kit. The protein concentration was normalized to 1-2mg/ml with PBS with 0.1% Triton X 100 or 1% SDS in PBS. 15µl of the lysates were loaded into each lane after mixed with 4xsample buffer+DTT and warmed at 42°C for 10min. Soluble and insoluble fractions were collected. Equal amounts of each fraction were separated by SDS-PAGE and transferred to PVDF membranes. After being blocked, blots were probed with Munc18-1 (D406V) Rabbit mAb #13414 (1:1000, Cell Signaling Technology) or Monoclonal ANTI-FLAG® M2 antibody produced in mouse (1:1000, F1804, MilliporeSigma). Blots were stained with Revert™ 700 Total Protein Stain for Western Blot Normalization (926-11011, LI-COR Technology) to confirm equal loading.

#### STXBP1 stability assay

HEK293T cells in 60 mm Petri dishes were transfected with different amounts of plasmid DNA to ensure a similar protein abundance, according to the plasmid DNA expression potentecy. 40 hrs post-transfection, cells were then treated with 10 µg/ml cycloheximide or vehicle control for 8 hours. Cells were lysed using RIPA buffer containing protease inhibitors. Equal amounts of each fraction were separated by SDS-PAGE and transferred to PVDF membranes. After being blocked, blots were probed with Munc18-1 (D406V) Rabbit mAb #13414 (1:1000, Cell Signaling Technology) or Monoclonal ANTI-FLAG® M2 antibody produced in mouse (1:1000, F1804, MilliporeSigma). Blots were stained with Revert™ 700 Total Protein Stain for Western Blot Normalization (926-11011, LI-COR technology) to confirm equal loading.

#### Co-immunoprecipitation of STXBP1 and STX1

TRE-T-SNARE stable cells were generated by transfection of HEK293T cells with 1µg of pPB-TRE-T-SNARE (pPB-TRE-STX1A-P2A-SNAP25-IRES-VAMP1) plasmid DNA+ 1µg of pPB-CAG TET-ON-mCherry plasmid DNA +1µg of pPB-CAG-PuroR plasmid DNA, 3µg of pPB-CAG-piggybac transposase plasmid DNA with DNA PolyJet™ In Vitro DNA Transfection Reagent (SL100688, SignaGen Laboratories). The transfected cells were selected with 1ug/ml puromycin (ant-pr-1, InvivoGen) for three days. The selected cells were growing to form clones in the plates. The clones were picked and propagated in 24 well plates. The T-SNARE (STX1A, SNAP2, and VAMP1) protein abundance was verified with Western blotting.

TRE-T-SNARE cells in 6xwell plates were transfected with different amounts of plasmid DNA to ensure a similar protein abundance, according to the plasmid DNA expression potency. 8 hrs post-transfection, The T-SNARRE protein expression was induced with 1µg/ml Doxycycline (A4052, APExBIO Technology LLC) 40 hrs later, the cells were lysed with IGEPAL™ CA-630 lysis buffer (50 mM Tris-Cl (pH7.5), 10% Glycerol, 150 mM NaCl, 0.5% IGEPAL™ CA-630) containing Protease Inhibitor Cocktail (EDTA-Free, 100X in DMSO) (B14002, Selleck Chemicals LLC), Phosphatase Inhibitor Cocktail (2 Tubes, 100X) (B15002, Selleck Chemicals LLC), and 10 mM 2-mercaptoethanol, followed by immunoprecipitation using anti-FLAG (B23102; Selleck Chemicals LLC). Following SDS-PAGE and transfer to PVDF membrane, samples were probed with anti-STX1 antibody (1:1000, 18572, Cell Signaling Technology).

### Principal component analysis and k-means clustering of functional data

Principal components were calculated from functional data on all variants. PC1 and PC2 accounted for most of the variation in the dataset (60.9% and 21.2%, respectively). PC1 is correlated with all four individual functional measurements; we use PC1 as a metric representing a combined functional score. K-means clustering was performed on principal components with two clusters specified (K=2).

## RESULTS

### EpiPred-STXBP1 accurately distinguishes between pathogenic and benign variants

We used our curated dataset of STXBP1 missense variants, which includes 80 BLB and 96 PLP variants annotated with 42 distinct features, to develop the single-gene classifier, EpiPred-STXBP1 (**Table S1, S2, S3**). We excluded any global VEPs that were trained on ClinVar data given that this was a primary resource for variants used to train EpiPred. We built a random forest model, which performed as a robust gene-specific classifier for our TS (**Figure S1**). Despite being trained on an substantially smaller dataset, EpiPred outperformed all global VEPs with an AUROC value of 98%, with the next two closest competitors having AUROC values of 90% (Varity) and 92% (ClinPred) (**Figure 1B**, **Table S4**)(*21, 33*). Indeed, EpiPred outperformed the global VEPs on sensitivity (96%), specificity (96%), and accuracy (96%) (**Table S4, Figure S2**). Several global VEPs that were not trained on ClinVar data, including AlphaMissense and VEST4, were prevalent features used in EpiPred, as assessed by SHAP, a game theory-based method to reveal feature impact on model decisions (*34*) (**Figure S3**). Other features that impact model predictions ranged from evolutionary conservation (PhyloP; 470way mammalian), residue solvent accessibility (RSA), and *in silico* prediction of protein stability (MAESTRO). These latter features strongly suggested that stability may be a key defining feature of *STXBP1* PLP missense variants, aligning with the decreased abundance and stability observed for missense variants in animal and cell models (*14–17*).

We used this model to score all variants with a Probability of PLP (EpiPred-P^PLP^ score) (**Figure 1C, D, Table S1**). Using a Gaussian mixture model, we defined the PLP threshold as an EpiPred-P^PLP^ score greater than 0.871 and the BLB threshold as an EpiPred-P^PLP^ score less than 0.625 (**Figure 1C**). In general, gnomAD variants present greater than thrice in the dataset had EpiPred-P^PLP^ scores below the PLP threshold (77/80 – 96% accuracy). The three exceptions were Phe153Ser and Arg536His, both predicted PLP and Ala553Thr in the ambiguous range (**Table S1, Figure S4)**. In the PLP TS 81% (78/96) variants exceeded the EpiPred PLP threshold, with nine variants in the ambiguous range and the remaining 10 PLP variants more likely to be benign by EpiPred (**Table S1, Figure S4)**. VUS (n = 150) values were scored broadly across the range, with 23% (35/150) in the PLP range and 55% (83/150) in the BLB range (**Figure 1C, D, Table S2**). We mapped these conflicting variants (10 PLP, 5 BLB) onto the 3D structure of the protein and assessed the SHAP outputs for each variant, but we were unable to detect any common features that could consistently explain this misclassification (**Figure S4**, **Figure S5**).

### Functional analysis of missense variants aligns with EpiPred predictions

We selected 20 variants for functional validation of EpiPred performance, including 10 BLB, 5 PLP, and 5 VUS. Of these VUS, three (p.Ala202Val, p.Trp288Leu and p.Ile576Thr) were predicted PLP by EpiPred and two (p.Gln301Leu, p.Lys333Glu) were predicted BLB by EpiPred (**Figure 1C, Table S5**). We used four functional readouts of variant pathogenicity that had been previously reported in models of *STXBP1* pathogenic missense variants. These included STXBP1 abundance (**Figure 2A**), solubility (**Figure 2B**), stability (**Figure 2C**), and the protein-protein interaction (PPI) between STXBP1 and STX1 (**Figure 2D**) (*12, 13*). Individually, all of these assays differentiated pathogenic from benign variants to some extent (**Table S6, S7, Figure S6-S9**). In particular the best discriminator was abundance; all of the PLP variants (n=5) showed statistically significant reduced abundance, with a mean reduction to 14% (range 4-42%) of wildtype levels, while only 5/10 BLB variants showed a significant change, with four variants reduced to ∼71% (range 50-73%) of wildtype, and one with 28% more protein than wildtype (**Figure 2A**). Similarly, protein stability was a good predictor of pathogenicity; all five PLP variants had statistically significant reduced stability (mean: 55%, range 38-64%), compared to only three of the BLB variants (n=3/10) with a smaller reduction (mean 80%, range 75-81%) (**Figure 2C**).

**Figure 2:**
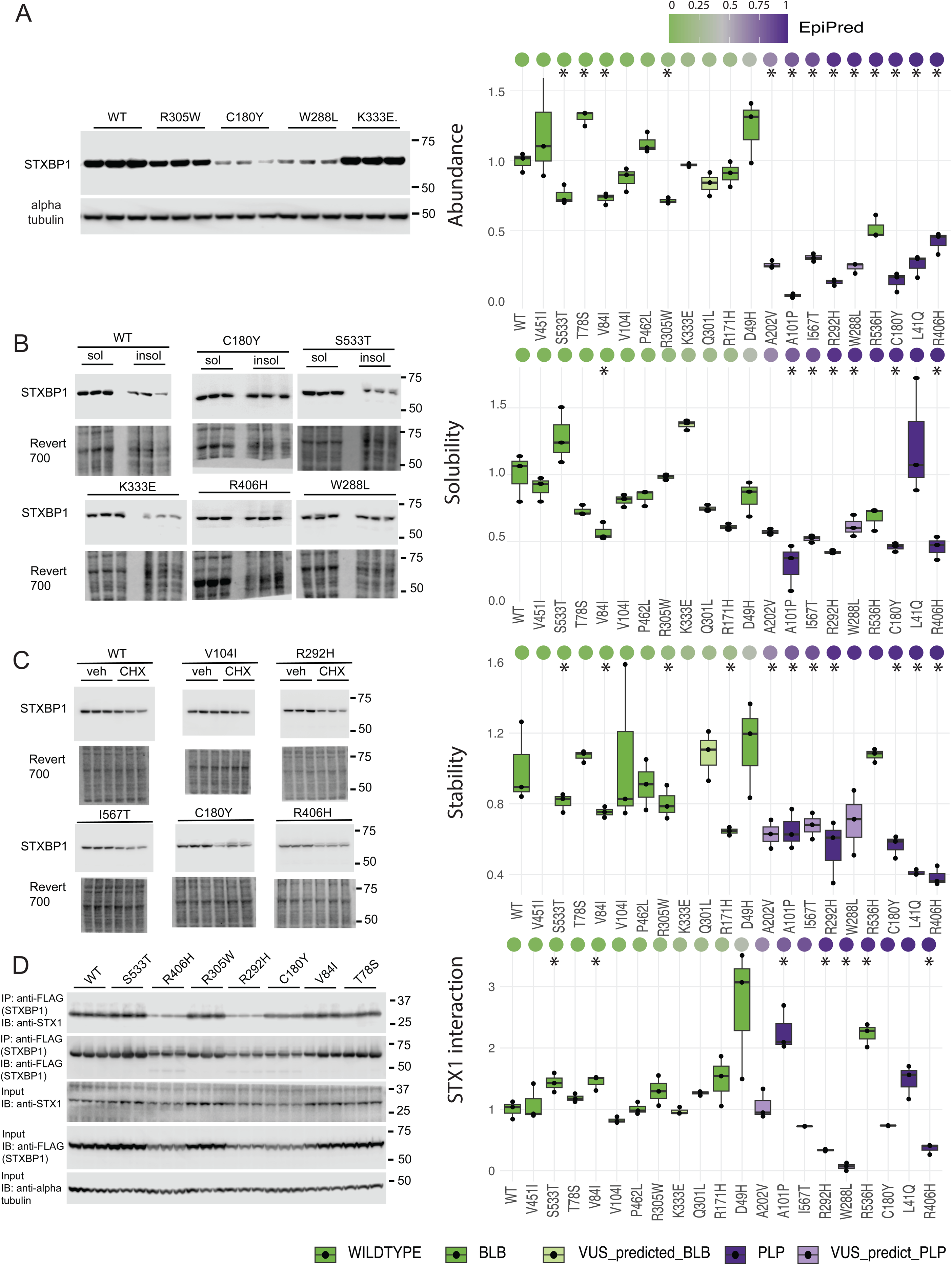
Biochemical assays and syntaxin 1 interaction distinguish pathogenic from benign STXBP1 missense variants. Data from representative western blots (left) and quantification (right) are plotted for the WT STXBP1 and twenty missense variants. Each assay is plotted separately: (A) Abundance, (**B**) Stability, (**C**) Solubility, and (**D**) STX1 PPI. Scores are normalized to the WT reference allele score. The quantified data is sorted, left to right, by EpiPred-P^PLP^ score from lowest (likely benign) to highest (likely pathogenic) and colored by class (wildtype, PLP, BLB, VUS predicted PLP/BLB). n=3, * p-value < 0.05, all raw data and p-values in **Table S5,6**.

There was a statistically significant correlation between stability and abundance (R=0.73, p=0.0005), as expected given the relatedness of these two assays (**Figure S10, S11**). All but one of the PLP variants (p.Leu41Gln) were less soluble (mean 41%, range 32-45%), compared to only one of the BLB variants (p.Val84Ile, 56% of wildtype). The STX1:STXBP1 interactions were more complex, 2 of the PLP variants (p.Arg292His and p.Arg406His) had statistically significantly reduced interactions (33% and 36% respectively); similarly, the predicted PLP variant p.Trp288Leu, interacted with STX1 at only 6% of wildtype levels. However, one PLP variant (p.Ala101Pro showed a > 2-fold increased interaction with STX1, as did three BLB variants, with a mean increase of 1.7-fold compared to wildtype (range 1.4-2.2). interaction. Overall, our VUS predictions by EpiPred were correct and the VUS that were predicted as either PLP or BLB clustered with known variants of each class.

In addition to these individual correlations, we considered the effects of the four functional readouts in combination and generated a combined functional score using principal component analysis (**Table S6)**. The first principal component accounted for 56% of the total variance and effectively distinguished between a BLB and a PLP cluster of variants (**Figure 3A**). All functional assays contributed to this distinction, with stability, and abundance the largest contributors, and STX1 interaction the least. Moreover, EpiPred had the strongest correlation with the combined functional score (0.72 compared to all other features, including global VEPs trained on ClinVar that were not used for EpiPred (**Figure 3B, S10, S12, Table S7**). Using this combined functional score, even the p.Arg536His variant clustered with the benign variants. This BLB variant, present in gnomAD (n=30), was predicted as pathogenic by EpiPred, and showed reduced abundance levels similar to PLP variants, with minimal effects on solubility and stability (**Figure 2**). However, p.Arg536His also demonstrated a >2 fold increased interaction with STX1, which led to a combined functional score in the BLB range. This example highlights the importance of leveraging multiple assays as a readout of missense impact on STXBP1 protein function, as the detrimental impacts of a missense variant in one assay may be ‘rescued’ by a different property of that variant in another.

**Figure 3:**
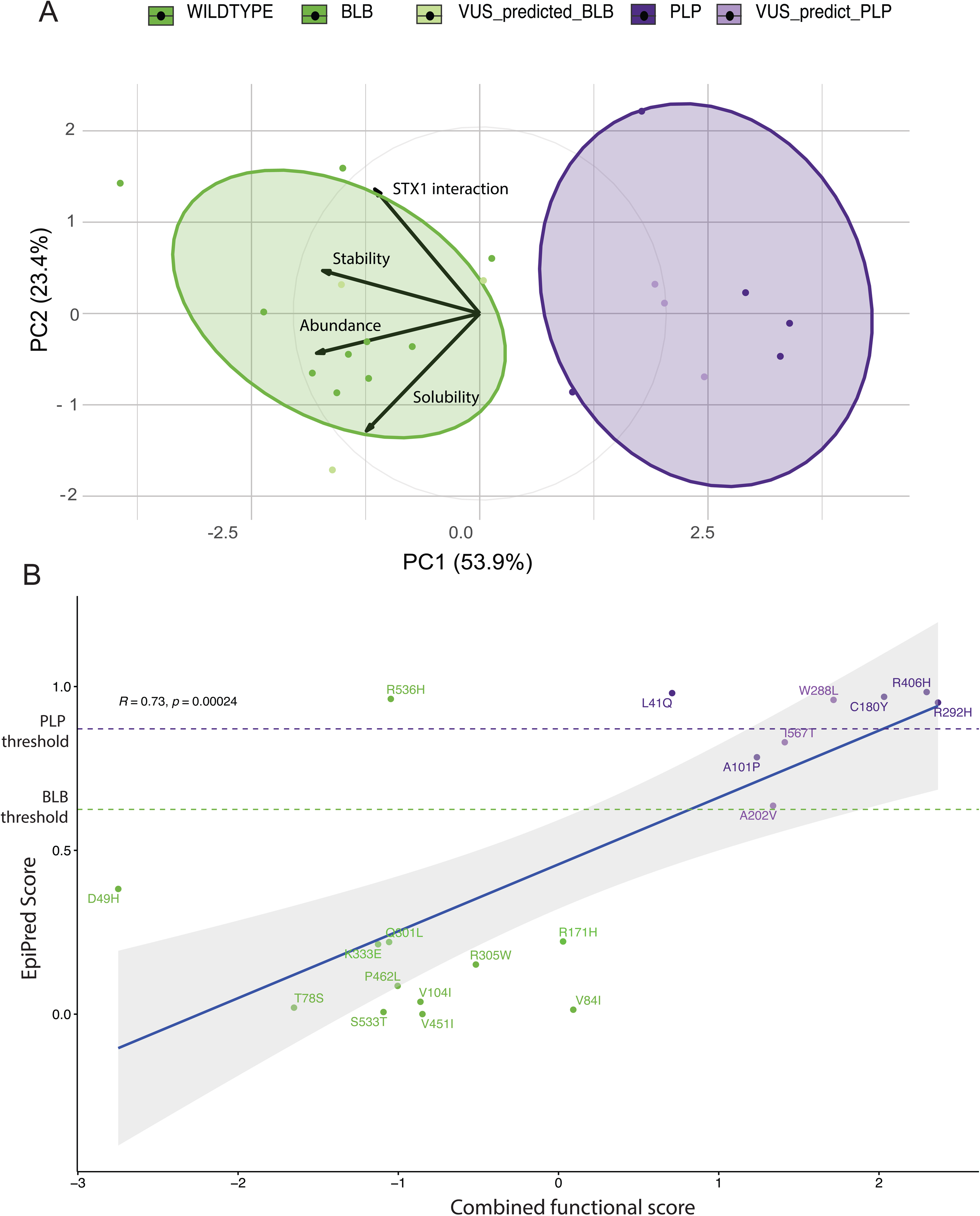
Integrating four functional assays with principal component analysis. (**A**) Biplot showing distinct BLB and PLP clusters within the dataset across two principal components, with PC1 explaining most of the variance in the data. (**B**) Scatterplot of EpiPred-P^PLP^ scores and combined functional score (PC1). EpiPred-P^PLP^ scores are highly correlated with the combined functional score. Variant and colored by class (PLP – dark purple, BLB – dark green, VUS predicted PLP-light purple, VUS predicted BLB, light green). (r=0.73; p=0.00024 Pearson).

### Evaluation of EpiPred in a holdout dataset demonstrates ability to detect misclassified disease-relevant missense variants

Leveraging the rapid pace of sequencing in population cohorts and individuals with epilepsy, we were able to establish a HS dataset consisting of 43 unique PLP missense variants, primarily from a large *STXBP1* phenotyping study(*30*), as well as 78 unique missense BLB variants (Regeneron exome server and gnomAD v4) (**Table S8**)(*35*). EpiPred, and indeed all VEPs investigated, exhibited reduced performance on the HS relative to the TS (**Figure 4A, S13-S17, Table S9**). All top performers in the test dataset, EpiPred (98%), Varity (90%) and ClinPred (92%), had reduced AUROCs in the HS: EpiPred (88%), Varity (87%) and ClinPred (80%) (**Table S9**). To explore this reduced performance, we evaluated the accuracy of the model for PLP and BLB variants in EpiPred. In the BLB HS EpiPred had a 92% (72/78) accuracy, with 3 variants predicted ambiguous and 3 predicted PLP, thus performing similarly to the TS (96% accuracy). In contrast, In the PLP HS accuracy diminished from 96% accuracy in the TS, to 67% (29/43) in the HS. This included five variants in the ambiguous range and the remaining 9 PLP variants predicted more likely to be benign by EpiPred (**Figure 4B)**. Thus, these findings revealed a potential misclassification of PLP variants in this cohort study. The criteria for PLP designation were described in the publication as “a modified annotation pipeline and only included variants that were either pathogenic or likely pathogenic by ACMG criteria or *de novo* variants in the coding region of STXBP1”. Moreover, we found good agreement between EpiPred and ClinPred/Varity predictions, with both global VEPs predicting ‘PLP’ missense variants as benign (n=8) or ‘ambiguous’ (n=1) (**Figure S15, S16, Table S10**). We re-evaluated the classification of these variants and determined most would not meet the ACMG criteria for PLP classification; indeed two (p.Asn398Ser and p.Ser253Asn) are classified as VUS in ClinVar. More specifically, the inheritance of these variants was not reported, three were found in individuals without epilepsy or in atypical phenotypes (e.g. focal cortical dysplasia), and these amino acid locations (though different substitutions) were present in the general population (gnomAD). Finally, two variants p.Asp207Gly and p.Asn398Lys showed no change in any of the biochemical assays or STX1 interactions that were used to assess the core STXBP1 phenotype (**Figure 4D, Table S5**). We thus reclassified these 9 variants as VUS and reassessed EpiPred classification performance with 34 PLP (from n=43 originally) and the same 78 BLB variants and observed increased AUROC performance in both EpiPred (97.8%) and global VEPs, Varity (95%) and ClinPred (93%) (**Figure 4C, S14 Table S11**). Both EpiPred and global VEPs (ClinPred/Varity) were still marginally outperformed by AlphaMissense (AUROC – 98.3%). This result illustrates the superior generalizability of AlphaMissense, but EpiPred still has greater sensitivity, which is better aligned with the goals of this VEP to identify individuals with pathogenic variants, i.e. true positives. Collectively, while the HS dataset had lower performance than the test set, EpiPred generalized well and still outperformed global VEPs. This analysis also highlights the power of EpiPred and global VEPs to accurately classify variants and identify false positives (and potentially misdiagnoses) in the literature.

**Figure 4:**
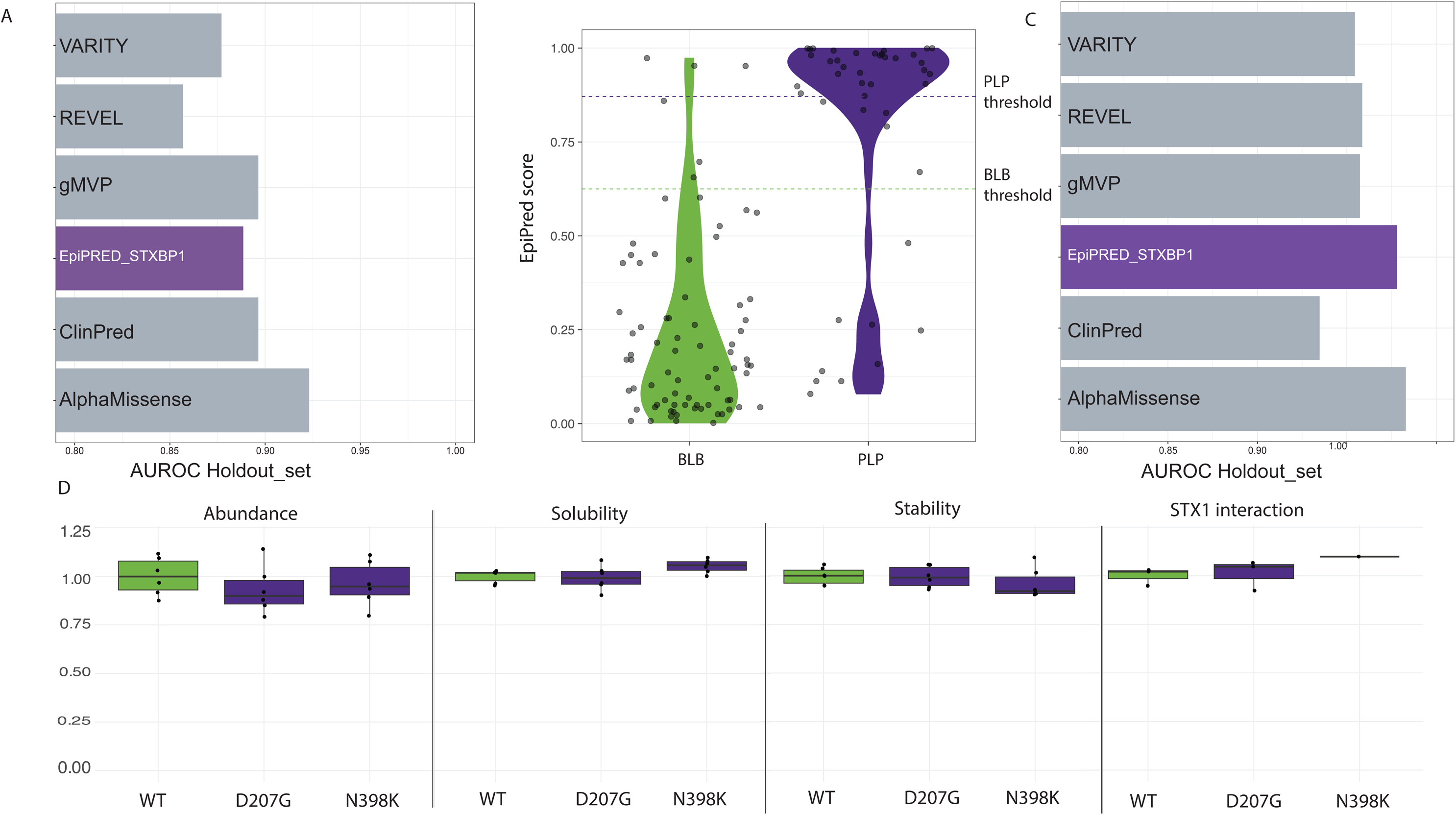
EpiPred classification of a holdout dataset. (**A**) Comparison of area under the ROC curve (AUROC) for EpiPred and a subset of high-performing globally-trained VEPs across the holdout dataset demonstrates reduced performance of EpiPred. (**B**) EpiPred-P^PLP^ score distributions in the full holdout data set shows the conserved performance for BLB variants, but a larger percentage of PLP variants with EpiPred-P^PLP^ scores in the benign/neutral range. EpiPred-P^PLP^ scores of 1 suggest pathogenicity, while scores of 0 suggest neutrality (**C**) Comparison of area under the ROC curve (AUROC) for EpiPred and a subset of high-performing globally-trained VEPs across the holdout dataset with reclassified variants, including the exclusion of 9 VUS from the PLP set, demonstrates improved EpiPred performance, on par with AlphaMissense. (**D**) Modeling of two misclassified variants, reclassified to VUS, which demonstrate properties similar to wildtype for abundance, stability, solubility and syntaxin interaction. n=6, * p-value < 0.05, all raw data and p-values in **Table S5**.

### Gene-wide EpiPred simulation reveals specific regions susceptible to perturbation

To facilitate dissemination of EpiPred to the scientific and epilepsy community we generated all possible missense *STXBP1* variants (n=3939) resulting from a single nucleotide change (**Table S12, Figure 5A**). We also collated data from the literature that functionally characterized missense variants (n=26) using both cellular and animal model systems and generated an aggregate functional score for each variant (**Table S13**)(*12–17*). This functional score, essentially a measure of how often a tested missense variant exhibited a phenotype, correlated reasonably well with EpiPred (**Figure S18**). Moreover, predicting STXBP1-related disorder (PRESR) was previously developed using functional data from cell-free assays and cellular models(*15*). We found that PRESR performed well, albeit not as well as EpiPred, for determining pathogenicity in both the TS and the HS (**Figure S19**). PRESR and EpiPred scores were highly correlated across all missense variants scored by both algorithms (r=0.77; Pearson), suggesting these *in silico* models may weight underlying features similarly (**Figure S20**). We next investigated the agreement between PRESR and our functional data. In general, PRESR predictions agreed with our functional results (18/20 variants; 90%). Both PRESR and EpiPred incorrectly predicted p.Arg536His (classified BLB) as pathogenic; PRESR further incorrectly predicted p.Asp49His (classified BLB) as pathogenic, while EpiPred correctly predicted p.Asp49His as benign (**Figure S20**). EpiPred more strongly correlated with the abundance assay and the combined score compared to PRESR (**Table S14**). Overall, our analyses suggest EpiPred slightly outperforms PRESR as a gene-specific classifier for *in silico* pathogenicity prediction of *STXBP1* missense variants.

**Figure 5:**
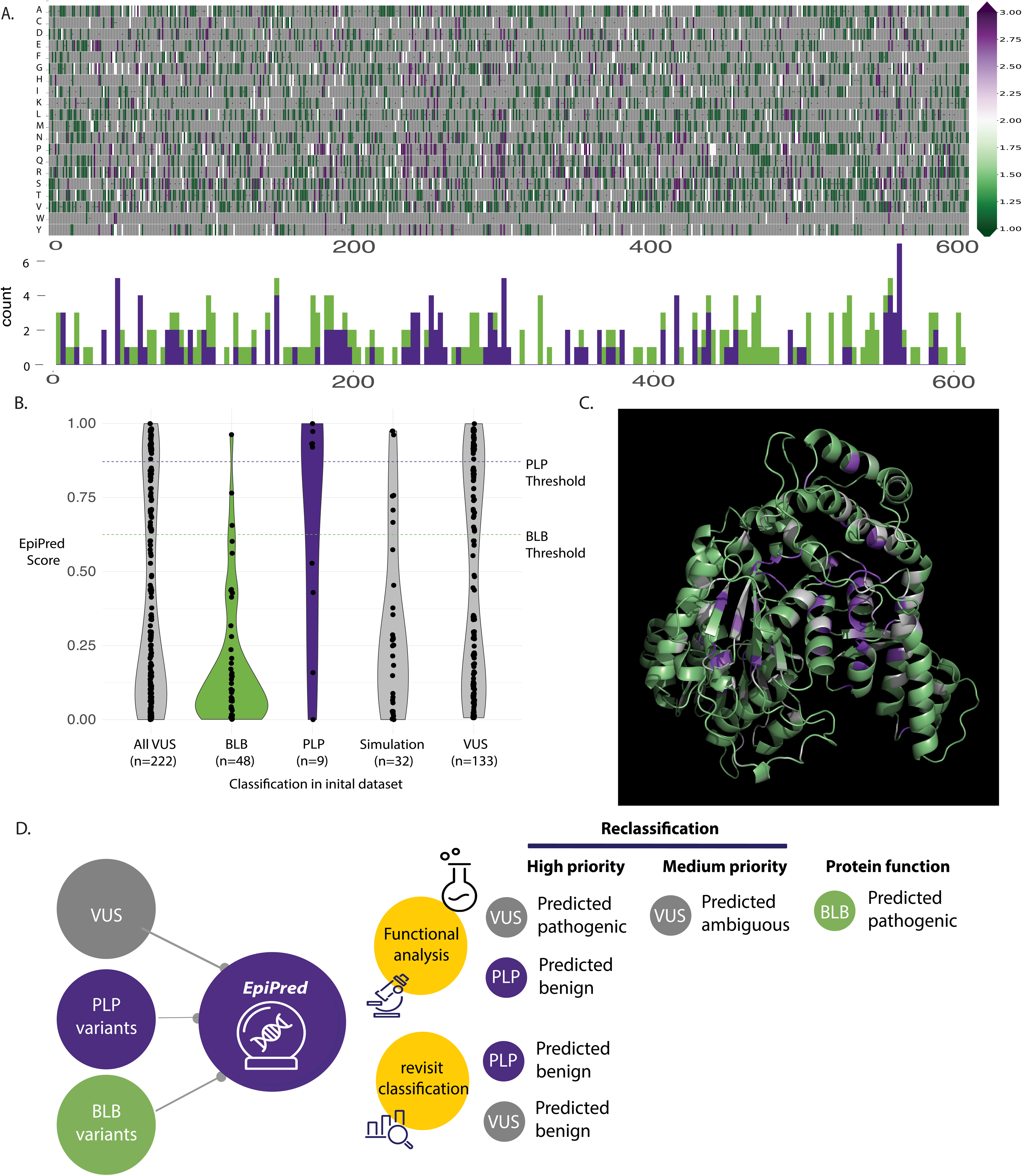
EpiPred-P^PLP^ scores for every possible STXBP1 missense variant and comparison with published functional data. **(A)** EpiPred-P^PLP^ scores for every possible SNV in the STXBP1 gene represented as a heatmap (top). Distribution of all known BLB and PLP variants across STXBP1 (bottom) (**B**) Distribution of EpiPred scores for latest version of ClinVar (accessed 8-7-2025) (**C**) EpiPred-P^PLP^ scores for every possible missense variant (mean per residue) were mapped onto the AlphaFold structure using PyMol. (**D**) Potential workflow for deploying EpiPred in reclassification of all classes of variants. For instance, those VUS EpiPred predicted as pathogenic, or PLP variants predicted benign are highest priority for functional interrogation to confirm pathogenicity. Conversely, those VUS and PLP variants predicted benign by EpiPred should be reevaluated in terms of the ACMG criteria for classification, as these may not be contributory.

Using our comprehensive STXBP1 EpiPred simulation, we scored all VUS in the latest ClinVar dataset, which now includes an expanded set of 222 variants, up from 150 when we initiated this study (**Figure 5B, Table S15, S16**). This includes 133 VUS from the original dataset, as well as 32 simulation or ‘new’ variants, which had comparable median EpiPred-P^PLP^ scores, 0.25 for new VUS, and 0.44 for previously identified VUS. There were, however, more VUS that were predicted to be pathogenic in the previously identified VUS (31/133 – 23%) as compared to the newly identified VUS (3/32 – 9%), though this difference was not statistically significant (Fisher’s Exact Test, Odds Ratio: 0.34, p-value of 0.092). There were also multiple VUS that were previously classified as BLB due to presence in gnomAD more than thrice, this group had a lower median EpiPred-P^PLP^ score (0.09). In addition, only 1 variant in this BLB group in predicted to be pathogenic (1/48 – 2%), which is far less than all VUS (40/222 – 18%) (Fisher’s Exact Test, Odds Ratio: 0.097, p-value of 0.0032). Finally, there were also 9 variants that were previously classified as PLP and have subsequently been reclassified as VUS. This includes four variants that were classified as benign by EpiPred, two variants that we reclassified from the HS (p.Asn398Ser, p.Ser253Ans) and two variants from the TS, p.Lys7Glu (EpiPred-P^PLP^ score 0.53) and p.Ser80Pro (EpiPred-P^PLP^ score 0.43) (**Figure S5, Table S10**). Overall, this reclassification analysis further demonstrates the utility of EpiPred to reclassify STXBP1 missense variants. To facilitate this, we developed a webapp, EpiPred (https://carvill-lab.shinyapps.io/epipred-shiny/) a Shiny-based user interface for clinicians, families and researchers to access our data and use the model to aid interpretation of STXBP1 missense VUS. We envision a workflow whereby EpiPred can assist in the reclassification of variants from clinical genetic testing, prioritizing high yield variants for functional characterization in the instance of predicted pathogenicity, and revisiting the ACMG classification criteria in the instance of predicted benignity (**Figure 5D**).

To visualize whether predicted pathogenic/benign variants cluster in specific regions of STXBP1, we aggregated EpiPred scores at each variant position and projected these onto the STXBP1 3D structure (**Figure 5C**). Rather than clustering within specific domains—which STXBP1 lacks—we found that known and predicted PLP variants tend to localize within the central core of the protein. This localization may underlie the reduced abundance and stability observed in STXBP1 proteins with pathogenic missense variants; highlighting a subset of variants, and patients, that could benefit from chaperone-based therapies such as PBA.

## DISCUSSION

Herein, we present EpiPred, a publicly accessible gene-specific classifier for pathogenicity prediction in the gene, *STXBP1*. EpiPred outperforms global VEPs, and our predictions were validated through four established functional assays that comprise a core *STXBP1* missense variant phenotype. This phenotype consists of reduced STXBP1 abundance, solubility, stability, and generally less interaction with syntaxin-1, though some PLP variants showed enhanced binding. While there was a strong correlation between EpiPred and both our and published functional data, there was a small subset of conflicting variants where our predictions did not match the known class. These variants represent a unique opportunity to understand protein function and the complex interaction of protein features, but also the ability to identify misclassified variants. We show that roughly a quarter of VUS could be pathogenic and develop an EpiPred Shiny app to facilitate the accessibility of variant interpretation and utility in clinical care. Beyond precision diagnostics, our computational and functional data consistently highlight reduced STXBP1 abundance as a key driver of pathogenic mechanisms. This insight paves the way for targeted precision therapies addressing protein abundance and instability, including promising approaches such as PBA-based treatments (*29*).

EpiPred outperforms all other global VEPs across multiple metrics, with an AUROC of 98%, as well as specificity, sensitivity and accuracy of 96% in the TS, with a 60:40 split of training and test datasets. These metrics were reduced in a modified HS dataset, which excluded nine VUS (see below), to an AUROC of 97.8%, specificity of 83%, sensitivity of 100% and accuracy of 88%. Even in the HS, EpiPred outperformed all global VEPs, with only AlphaMissense (AUROC 98.3%) slightly outperforming EpiPred in the holdout set; by comparison, the AUROC of AlphaMissense in the TS was 94%. Thus overall, EpiPred outperformed all other global VEPs, despite being trained on a substantially smaller dataset. This is in keeping with the handful of other gene-specific ensemble VEPs, particularly those for *BRCA1*/2, which demonstrated a 96% sensitivity and 98% sensitivity(*26*). Collectively, we demonstrate the utility of gene-specific VEPs for clinically relevant genes, and our goal in EpiMVP is to extend this approach to other non-ion channel related genes, prioritizing those more commonly implicated in epilepsy: *SYNGAP1*, *SLC6A1* and *PCDH19*.

We also propose a framework by which EpiPred could be deployed for reclassification of STXBP1 missense variants including prioritizing variants for functional validation. Specifically, VUS predicted to be pathogenic by EpiPred, as well as known pathogenic variants predicted to be benign, warrant closer examination. While EpiPred demonstrates strong predictive performance, it remains essential to functionally characterize variants to uncover how distinct features of STXBP1 missense variants interact and contribute to disease mechanisms. For instance, one of the most intriguing variants in this study was p.Arg536His classified as BLB and present in 30 individuals in gnomAD, primarily in individuals of non-Finnish European ancestry (n=27). However, EpiPred and indeed all other global VEPs predict p.Arg536His as likely pathogenic (EpiPred-P^PLP^ score = 0.96). In our heterologous system, this variant exhibited reduced abundance and solubility compared to wildtype, in keeping with the majority of the pathogenic variants. However, protein stability was unchanged, while the variant displayed a two-fold increased binding of STX1. Thus, one could speculate that increased STX1 binding might ‘rescue’ the reduced abundance of the mutant protein. In keeping with this logic, the combined functional score suggested neutrality, as the variant clustered with other BLB variants. This observation highlights the strength of evaluating multiple putative biochemical consequences of a missense variant and provides motivation for the combined functional score. Another key example is p.Ala101Pro, a PLP variant which also had ∼2-fold increase in STX1 interaction, but greatly reduced abundance, stability and solubility, which is presumably insufficient to offset the increased STX1 interaction. In this instance, both EpiPred and the combined functional score correctly predict this variant as pathogenic. We also speculate that a number of the conflicting variants may display similar complex interactions between the SNARE protein binding partners as well as the biophysical and biochemical properties of the mutant protein. Overall, these two examples, as well as the other fifteen discordant variants should be prioritized for functional follow-up, both within our system and in more pathologically relevant models, such as iPSC-derived neurons and animals. For instance, in our EpiMVP companion paper we modeled truncation (p.Gly544Val-fs*2) and missense p.Ty75Cys (EpiPred-P^PLP^ score = 0.98) variants derived from individuals with DEE in human iPSC-derived neural precursors and glutamatergic neurons. Similar to our overexpression data, we demonstrate decreased STXBP1 and STX1 protein levels, as well as decreased solubility. Glutamatergic neurons from these patients also exhibited reduced synaptogenesis, increased neurodegeneration, and decreased neuronal activity and network synchrony. Collectively, our studies implicate the core mutant STXBP1 protein phenotype as an integral molecular component of disease pathophysiology. A systematic analysis of conflicting variants offers a valuable opportunity to deepen our understanding of protein function and the intricate interplay of features underlying the core STXBP1 missense variant phenotype.

Our functional and computational analyses reveal that, rather than cluster in specific domains, which STXBP1 does not have, PLP variants tend to occur in the middle core of the protein. Further, RSA, a measure of how buried an amino acid residue is within a protein, was one of the primary drivers of EpiPred, suggesting the core of STXBP1 is more likely to harbor disease-related variants. This aligns with global protein predictions, where pathogenic variants generally tend to be more buried within a protein than polymorphisms that tend to be exposed(*36*). RSA is also a proxy for stability of a protein, which was one of the strongest functional discriminators of pathogenic and benign variants, again highlighting the strength of combining functional and *in silico* predictions. There are now multiple studies, including our companion EpiMVP paper, which show PBA can rescue phenotypes associated with STXBP1 loss in yeast, C.*elegans*, mouse neurons(*17*), and iNeurons (*37*). These studies are timely, as PBA is currently in clinical trials for *STXBP1*-related disorder(*29*). In our companion EpiMVP paper we show that two patients in this PBA trial, with a missense variant, p.Arg292Pro (EpiPred-P^PLP^ score = 0.98), and a truncating variant (p.Ser311PhefsX3) had reduced seizure frequencies post treatment(*37*). Collectively, our data suggest that individuals with missense variants that are confirmed or predicted to be unstable might benefit from PBA, or similar, therapies. While stability of mutant proteins was one of the biggest drivers of pathogenicity of EpiPred. In the future, high-throughput methods such as VAMP-seq, which uses GFP-tagged proteins and massively parallel sequencing to measure steady-state protein abundance, could be used to create a comprehensive map of STXBP1 missense variant abundance (*38*). This approach was used successfully to evaluate the entire coding sequence of *PTEN*(*39*), *CYP2C9*(*40*), *Parkin*(*41*), and other clinically relevant genes, revealing key domains prone to instability or targeted by proteostasis.

Further, for variants that are classified in ClinVar as pathogenic but predicted by EpiPred to be benign, there is a strong case that these variants may indeed be neutral. We propose a framework for deployment of EpiPred after initial classification by genetic testing. For those variants that EpiPred predicts as benign functional characterization is necessary, including multiple ‘PLP’ variants in the holdout dataset that are likely neutral. The vast majority of pathogenic variants in the holdout dataset were from a large cohort study (n=534) examining the clinical features of STXBP1-related NDD(*42*). We reclassified nine of these variants as VUS, using ACMG criteria and indeed two (p.Asn398Ser and p.Ser253Asn) are classified as VUS in ClinVar. These decisions were based on the following evidence: good agreement between EpiPred and other global VEPs that these variants are more likely neutral, the inheritance of these variants was not reported, and these amino acid locations (though different substitutions) were present in the general population (gnomAD). Functional studies for two of these variants (p.Asp207Gly and p.Asn398Lys) demonstrated no change in STXBP1 abundance, stability, solubility, or interaction with STX1, in keeping with wildtype STXBP1 function. Thus, collectively it is likely that some of these VUS, with the exception of p.Asp207Gly as discussed below, are misdiagnoses, and further functional characterization in our iNeuron system is warranted. In addition, three of these variants were identified in individuals without epilepsy, or in atypical phenotypes (e.g. focal cortical dysplasia); we speculate that these variants may indeed be pathogenic, just not for epilepsy, and that EpiPred, having been trained solely on epilepsy-related variants, may misclassify these variants. Again, further functional analyses, using MAVEs and iNeurons are needed to support this hypothesis. Moreover, these insights will be advanced as additional further genotype-phenotype correlations as more individuals with rare *STXBP1* variants are identified in individuals with neurological conditions without epilepsy.

Despite the high predictive power of EpiPred, it still has limitations. For instance, in the holdout dataset, the variant p.Asp207Gly was predicted as benign by all global VEPs and EpiPred (EpiPred-P^PLP^ score = 0.13) and had the same biochemical properties and syntaxin interaction as wildtype STXBP1 in functional studies. However, iPSCs derived from the patient carrying this variant revealed reduced mRNA and protein levels of STXBP1, possibly due to the generation of a cryptic splice site(*14*). Unfortunately, this aberrant splicing was not explored in the iPSC model, and an additional missense variant (p.Arg235Gln) showed reduced mRNA levels, suggesting that reduced mRNA is a more general feature of pathogenic missense variants that warrants further study. However, this p.Asp207Gly variant does highlight an important limitation of our study, that missense variants that disrupt splicing via alteration of an exonic splice enhancer cannot be detected in EpiPred or our heterologous overexpression-based system. The most popular splice-site prediction tool, SpliceAi, did not predict p.Asp207Gly to disrupt splicing; however, the misclassified PLP variant, p.Glu487Asp with a neutral EpiPred-P^PLP^ score (0.48) is predicted to disrupt an existing donor site and create a new one ∼100bp upstream. Thus, tools such as SpliceAi(*43*), and other predictors focused on motifs within exons, should be incorporated into EpiPred and other gene-specific predictors in the future.

To facilitate the utility of, and access to, EpiPred, we built a user-friendly Shiny-based web app that allows users to input their *STXBP1* missense variant of interest and obtain the EpiPred-P^PLP^ score. Furthermore, the user receives information to aid in interpretation, including an explainer of variant classes and what the prediction means, as well as where the variant lies in the 3D structure. Our intention is that the ‘Check my Variant’ tab be utilized by clinicians, genetic counselors, parents, and caregivers, to add interpretation of an STXBP1 missense variant, though all results should be considered in combination with other clinical and genetic data, preferably with a genetic counselor and the clinical care team. We also make the same information available in the ‘For Researchers’ tab, where EpiPred predictions for every possible missense variant can be compared to global VEP predictions, correlations between VEPs can be visualized, and other features including gnomAD allele counts and amino acid positions are delineated.

As part of the design of the EpiPred Shiny app, we generated every possible missense variant that could result from a single nucleotide change (n=3492 missense variants). Of these,13% (n=466) are predicted to be pathogenic by EpiPred. Moreover, of the 151 VUS in our original ClinVar dataset, 23% (n=35) have scores in the pathogenic range, thus suggesting that many existing and putative VUS may be related to the disease phenotype. While global VEPs and functional data are routinely used as ‘supporting’ and ‘supporting to moderate’ evidence in the ACMG framework, our study suggests that integrating EpiPred could have high clinical utility. In the current study, we propose a model whereby EpiPred is deployed after initial variant classification leveraging the webapp and community engagement. Future systematic incorporation of EpiPred for reclassification will most readily be achieved through the Epilepsy Gene Variant Curation Groups within the ClinGen initiative. Within this framework, it will be important to not only flag VUS that may be pathogenic to facilitate a genetic diagnosis but also examine those VUS that are currently misclassified as pathogenic. These insights are also integral given the emerging precision therapy opportunities for individuals with STXBP1-related NDD, including the clinical trial for PBA. Identification of patients with a precise genetic diagnosis is crucial for enrollment in a precision therapy trial. Finally, both our computational and functional analyses demonstrate abundance and stability as key features associated with *STXBP1* pathogenic missense variants. Coupled with VAMP-seq in the future, this feature will allow us to *a-priori* define those missense variants that perturb abundance, and thus the individuals that may best benefit from PBA or more targeted small molecule chaperones.

## Acknowledgments

This study was sponsored by the NIH EpiMVP CWOW NINDS U54 NS117170

