## Supplementary material for "EpiPred: A gene-specific machine learning model for classifying missense variants in the epilepsy-related gene *STXBP1*": Figure S1

**Supplemental Table 11: Comprehensive model metrics for each annotated feature across holdout set with updated Holdout Set minus variants reclassified to VUS (n=9).** **4**

**List of supplementary tables**

**Supplementary Table 1: All truth set STXBP1 variants annotated with EpiPred model output**
**Supplementary Table 2: All STXBP1 missense VUS annotated with EpiPred model output**
**Supplementary Table 3: Feature lists for EpiPred model training and benchmarking**
**Supplementary Table 4: Comprehensive model metrics for each annotated feature and collection of global VEPs across full truth set**
**Supplementary Table 5: Quantification of all western blot data for the 20 modeled missense variants**
**Supplementary Table 6: Statistics for all western blot data for the 20 modeled missense variants**
**Supplementary Table 7: Pearson correlation table to demonstrate correlation of each functional assay with each annotated feature**
**Supplemental Table 8: All holdout set STXBP1 variants annotated with EpiPred model output**
**Supplemental Table 9: Comprehensive model metrics for each annotated feature across holdout set.**
**Supplemental Table 10: PLP Variants called in the Truth and Holdout dataset predicted to be BLB by EpiPred**
**Supplemental Table 11: Comprehensive model metrics for each annotated feature across holdout set with updated Holdout Set minus variants reclassified to VUS (n=9).**
**Supplemental Table 12: All possible missense variants in STXBP1 along with metrics and EpiPred predictions**
**Supplemental Table 13: Published functional data for a subset of missense variants**
**Supplemental Table 14: Pearson and spearman correlation of PRESR model with functional data. EpiPred correlation also shown for comparison.**
**Supplemental Table 15: All VUS in current version of ClinVar (date accessed 8-4-2025)**
**Supplemental Table 16: Summary of EpiPred properties of all VUS in current version of ClinVar (date accessed 8-4-2025)**
Please see separate excel document

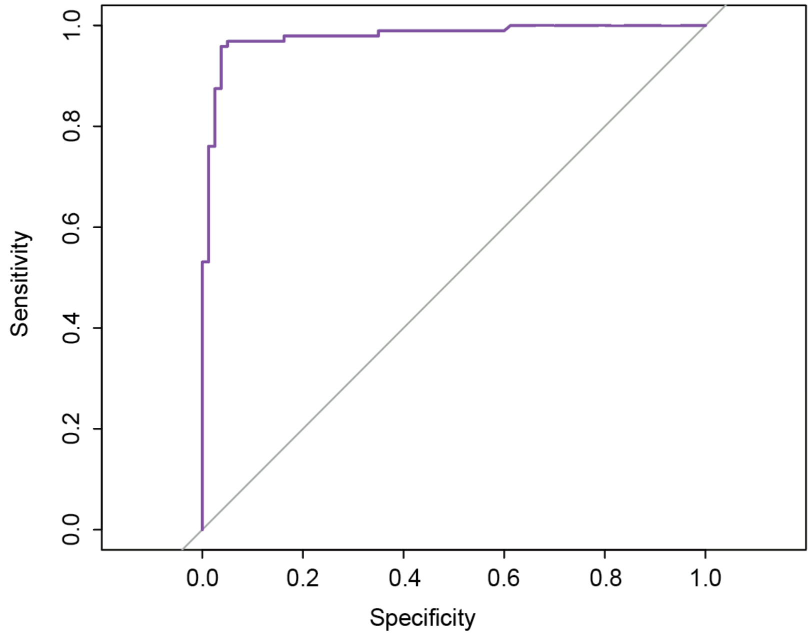

**Supplementary Figure 1: AUROC for the EpiPred truth set**

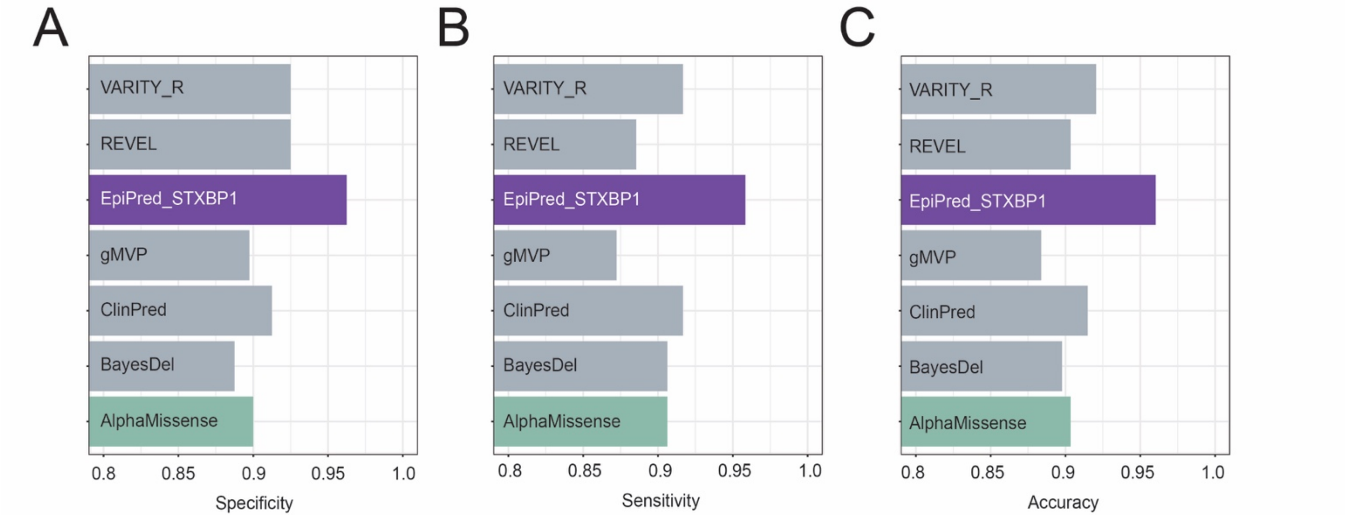

**Supplementary Figure 2: Specificity, sensitivity, and accuracy of EpiPred and other VEPs on the full truth set**

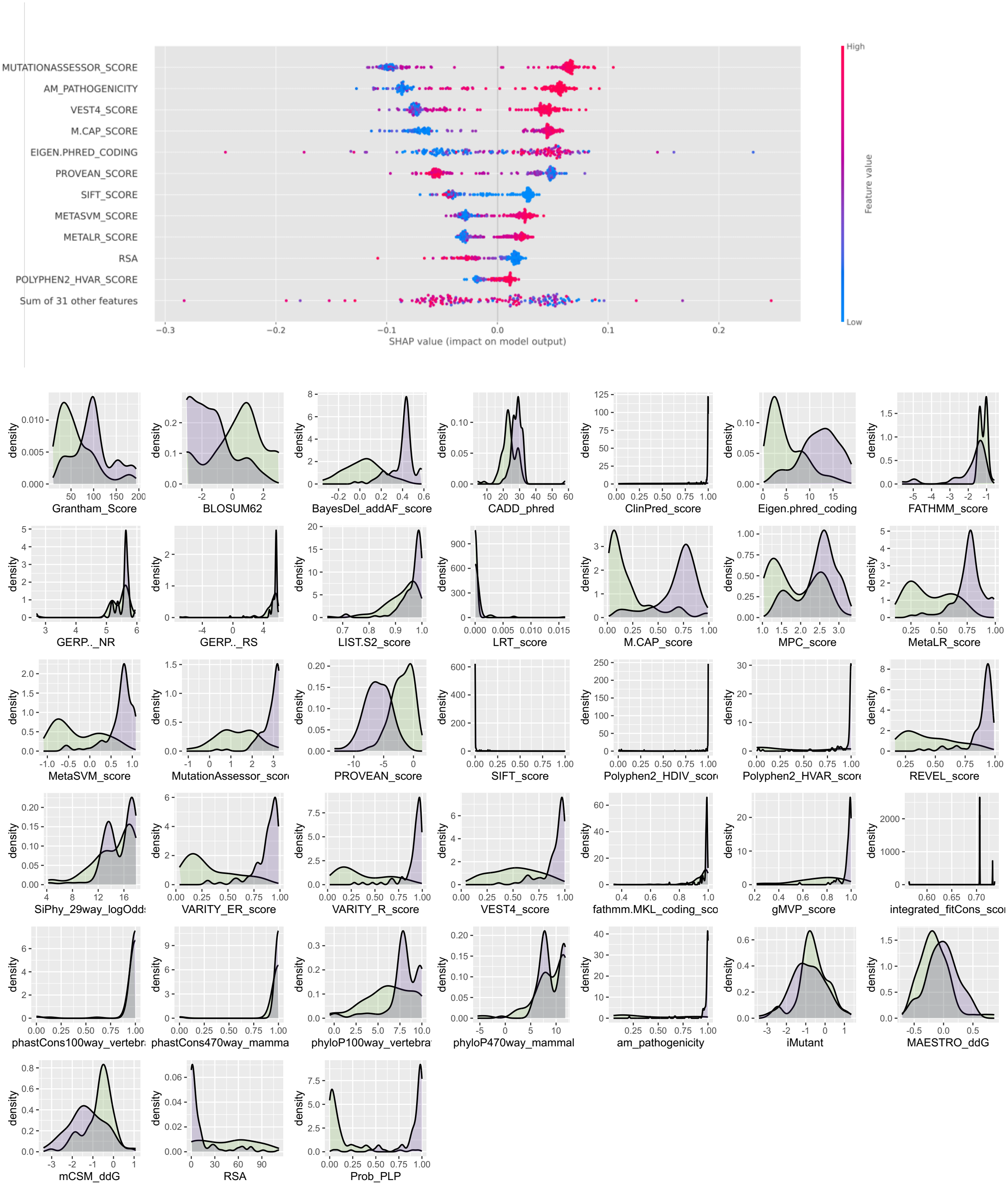

**Supplementary Figure 3: SHAP distributions of feature importance for the truth set (top).** The top features leveraged in EpiPred are the global VEPs, with Mutation Assessor being the strongest contributor, residue solvent accessibility (RSA) also ranks highly. Feature distribution for EpiPredSTXBP1 in the truth set (bottom). Green is class 0 (Truth set BLB); purple is class 1 (Truth set PLP). For unabbreviated feature names, see Table S3.

### BLB missense variants that are predicted pathogenic by EpiPred

Phe153Ser (gnomAD allele count 2)

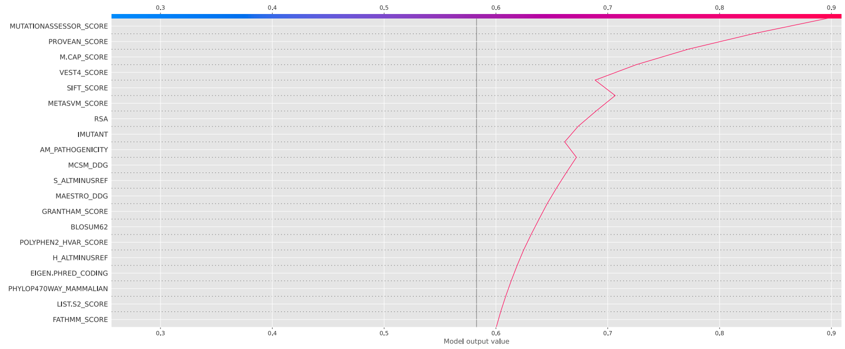

Arg536His (gnomAD allele count 30)

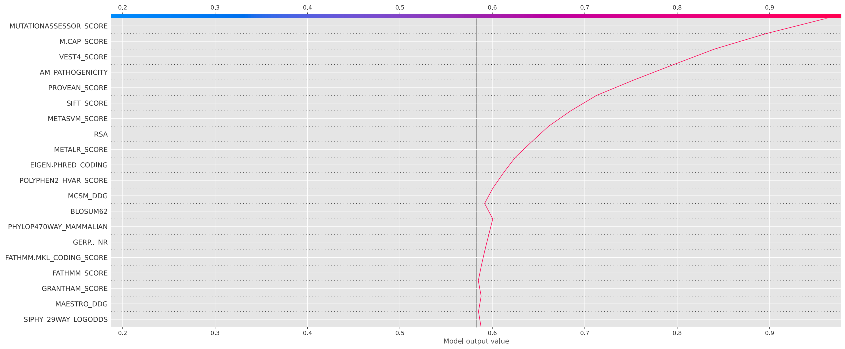

### PLP missense variants that are predicted benign by EpiPred

Gly193Val

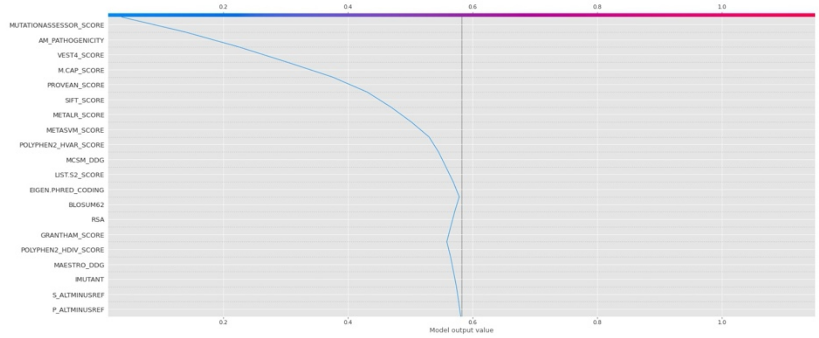

### Asn548Asp

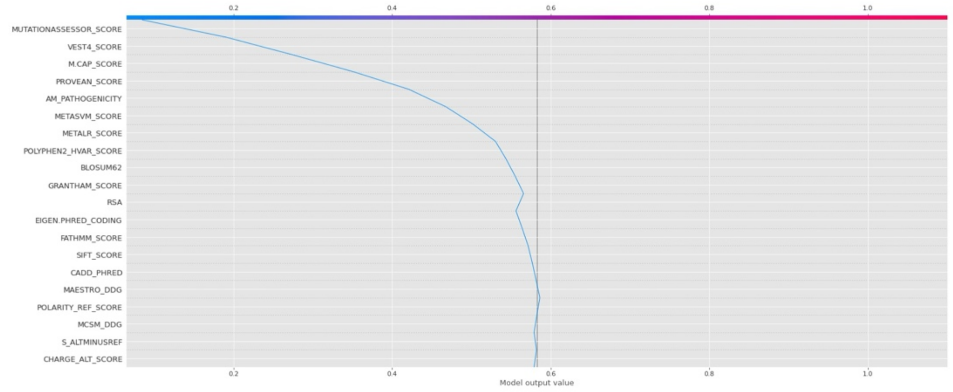

### Ala517Ser

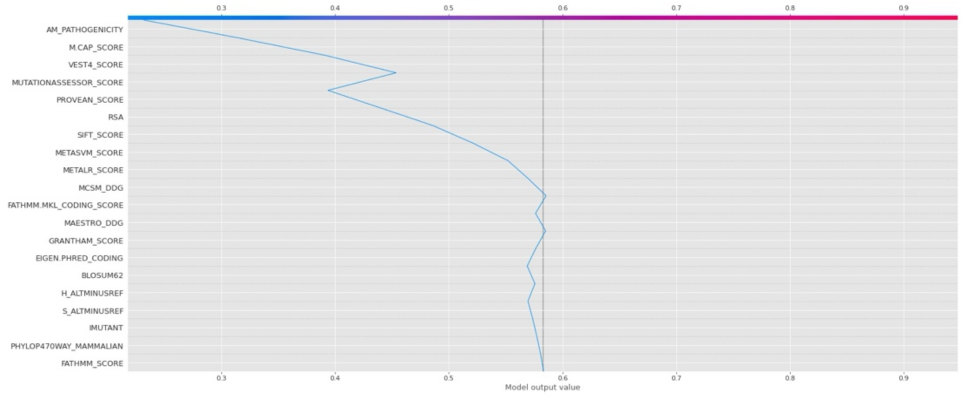

### Ser80Pro

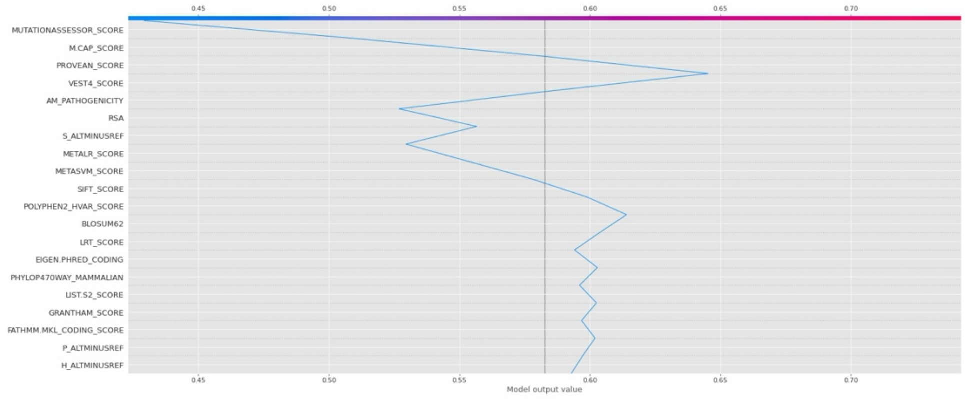

Glu487Asp

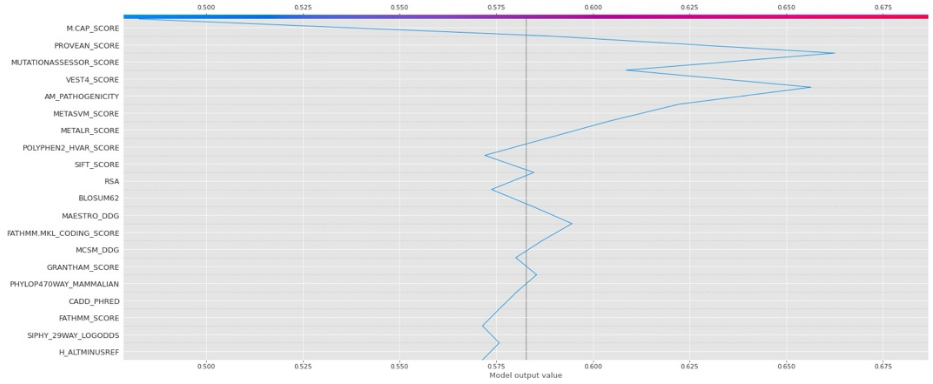

Thr129Pro

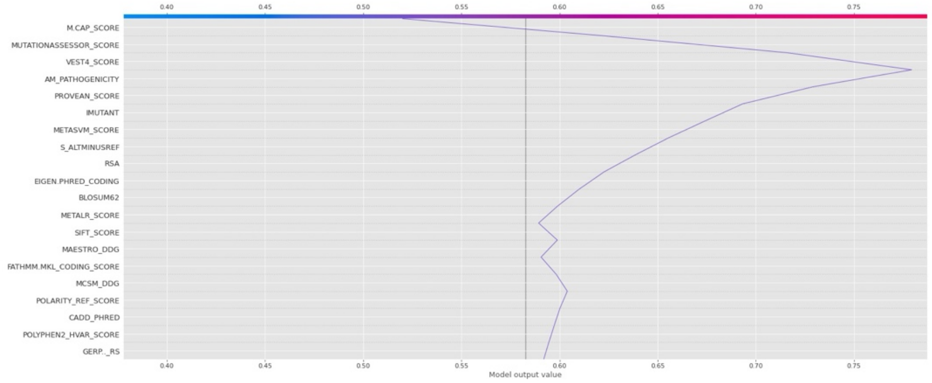

Lys7Glu

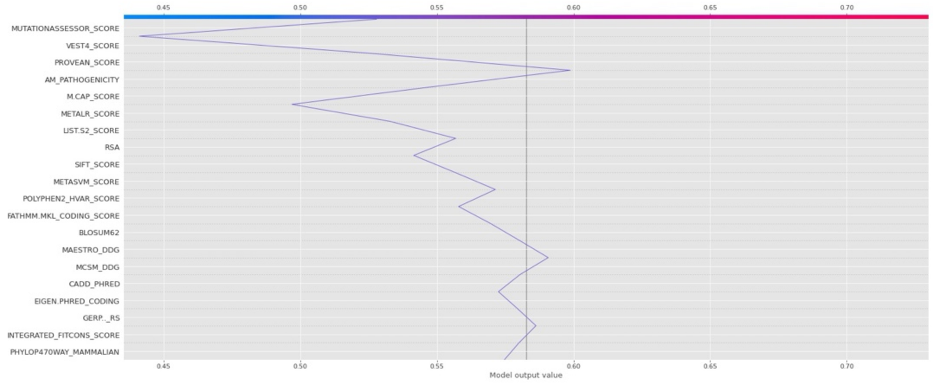

Asp284Tyr

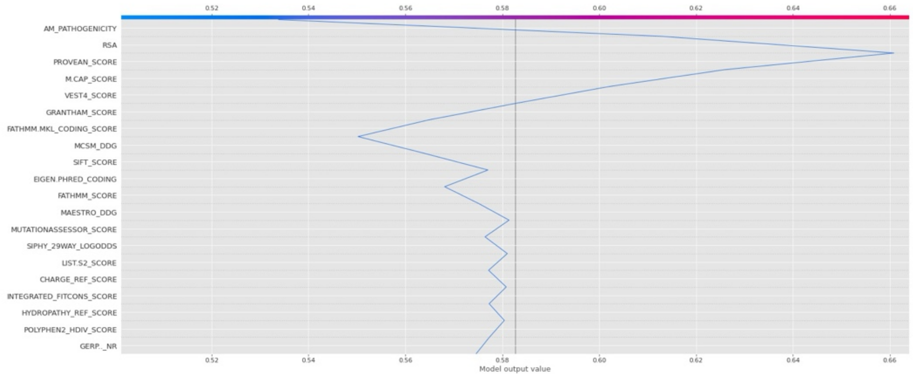

Thr574Pro

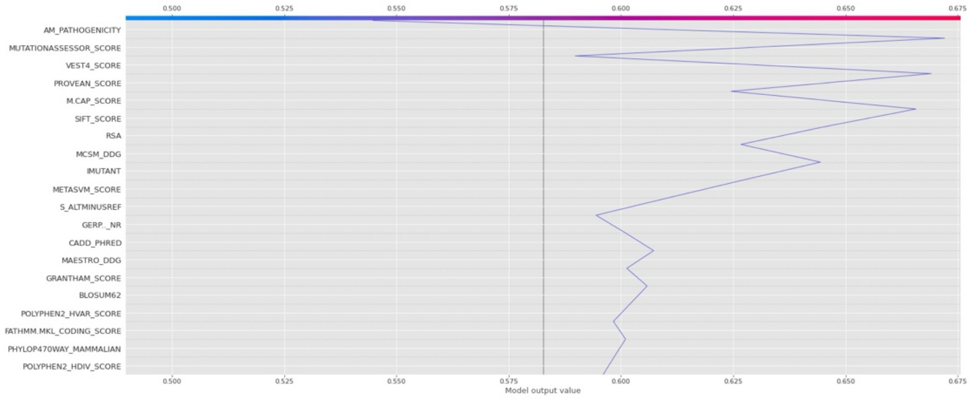

Thr361Ile

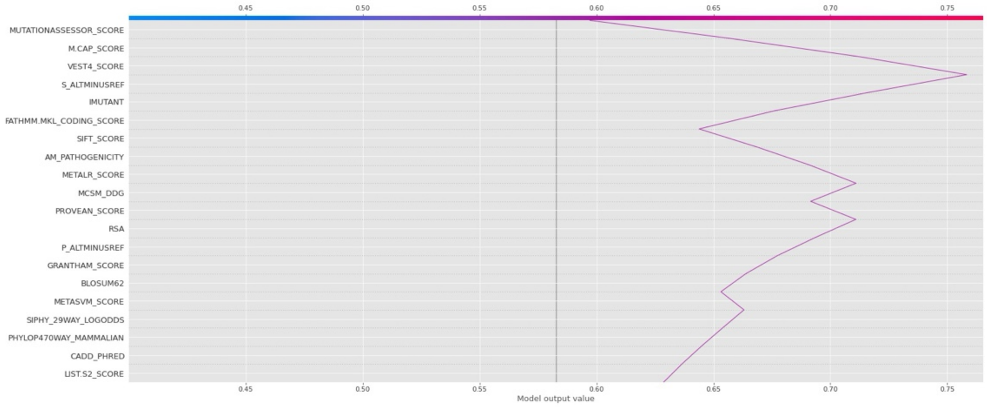

**Supplementary Figure 4: Variants from the truth set that are incorrectly called by EpiPred**

| Conflicting variants (n=15) in the truth set called the incorrect class by EpiPred-STXBP1 | | | | |
|---|---|---|---|---|
| **Variant** | **Class** | **EpiPred score (prediction)** | **SpliceAi score (max)*** | **Variant at same position** |
| K7E | PLP | 0.53 (BLB) | 0.01 | No |
| S80P | PLP | 0.43 (BLB) | 0 | No |
| T129P | PLP | 0.52 (BLB) | 0 | No |
| G193V | PLP | 0.04 (BLB) | 0.49 (donor loss) | No |
| D284Y | PLP | 0.53 (BLB) | 0.01 | No |
| T361I | PLP | 0.60 (BLB) | 0.01 | No |
| E487D | PLP | 0.48 (BLB) | **0.78 (donor loss)** | No |
| A517S | PLP | 0.23 (BLB) | 0 | Yes, A517T, BLB |
| N548D | PLP | 0.08 (BLB) | 0 | No, but also N548I in the holdout set PLP called BLB by EpiPred (0.26) |
| T574P | PLP | 0.54 (BLB) | 0.05 | No |
| H103D | BLB | 0.97 (PLP) | NA | Yes, H103P, PLP |
| F153S | BLB | 0.90 (PLP) | NA | No |
| Y266C | BLB | 0.95 (PLP) | NA | No |
| R536C | BLB | 0.95 (PLP) | NA | No |
| **R536H** | **BLB** | **0.96** (PLP) | NA | **No** |

***** SpliceAi gives four scores for acceptor/donor gain/loss we show only highest score, 0.8 indicates a site predicted likely to disrupt splicing

Variants in red classified as VUS in most recent version of ClinVar

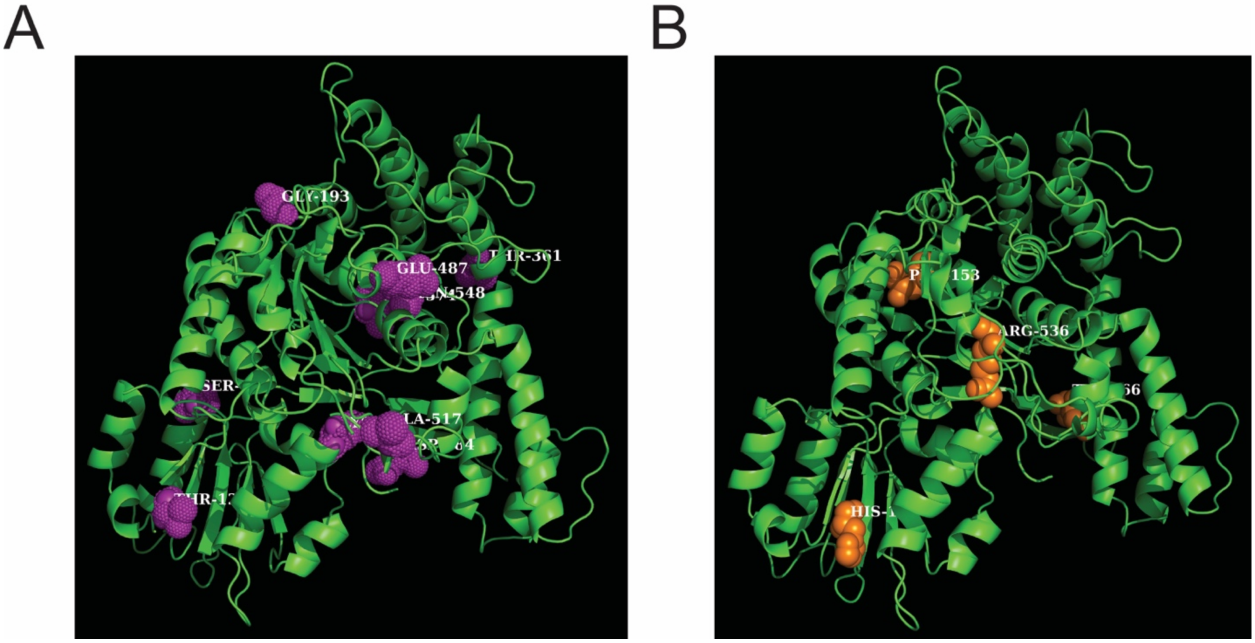

**Supplementary Figure 5: Location of conflicting variants in the 3D representation of STXBP1**

The 10 PLP variants (left in purple) and 5 BLB variants (right in orange) that are incorrectly predicted by EpiPred and other global variant effect predictors i.e. where the known ACMG class and EpiPred do not match.

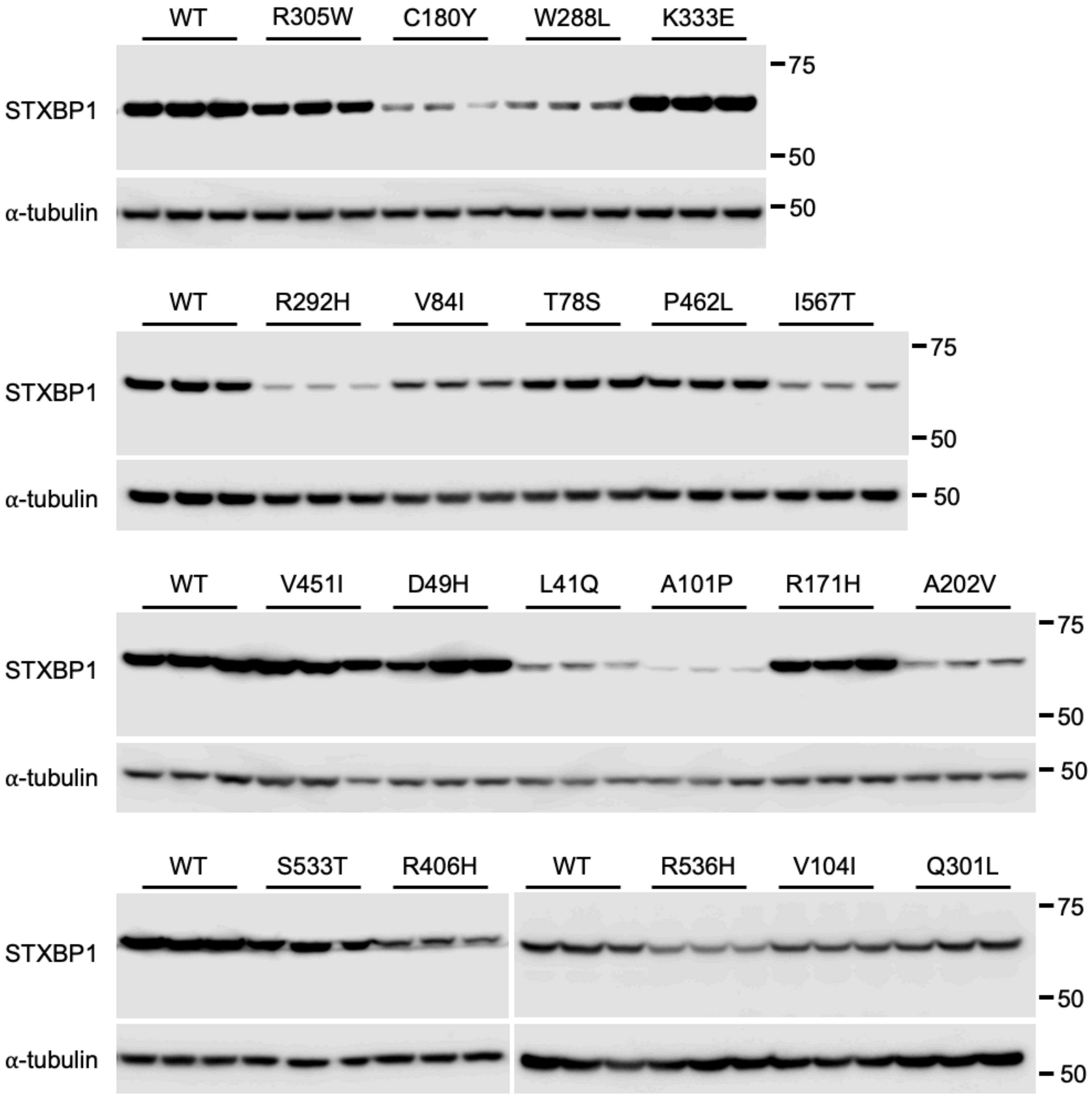

WT
R305W
C180Y
W288L
K333E
STXBP1
α-tubulin
75
50
50
WT
R292H
V84I
T78S
P462L
I567T
STXBP1
α-tubulin
75
50
50
WT
V451I
D49H
L41Q
A101P
R171H
A202V
STXBP1
α-tubulin
75
50
50
WT
S533T
R406H
WT
R536H
V104I
Q301L
STXBP1
α-tubulin
75
50
50

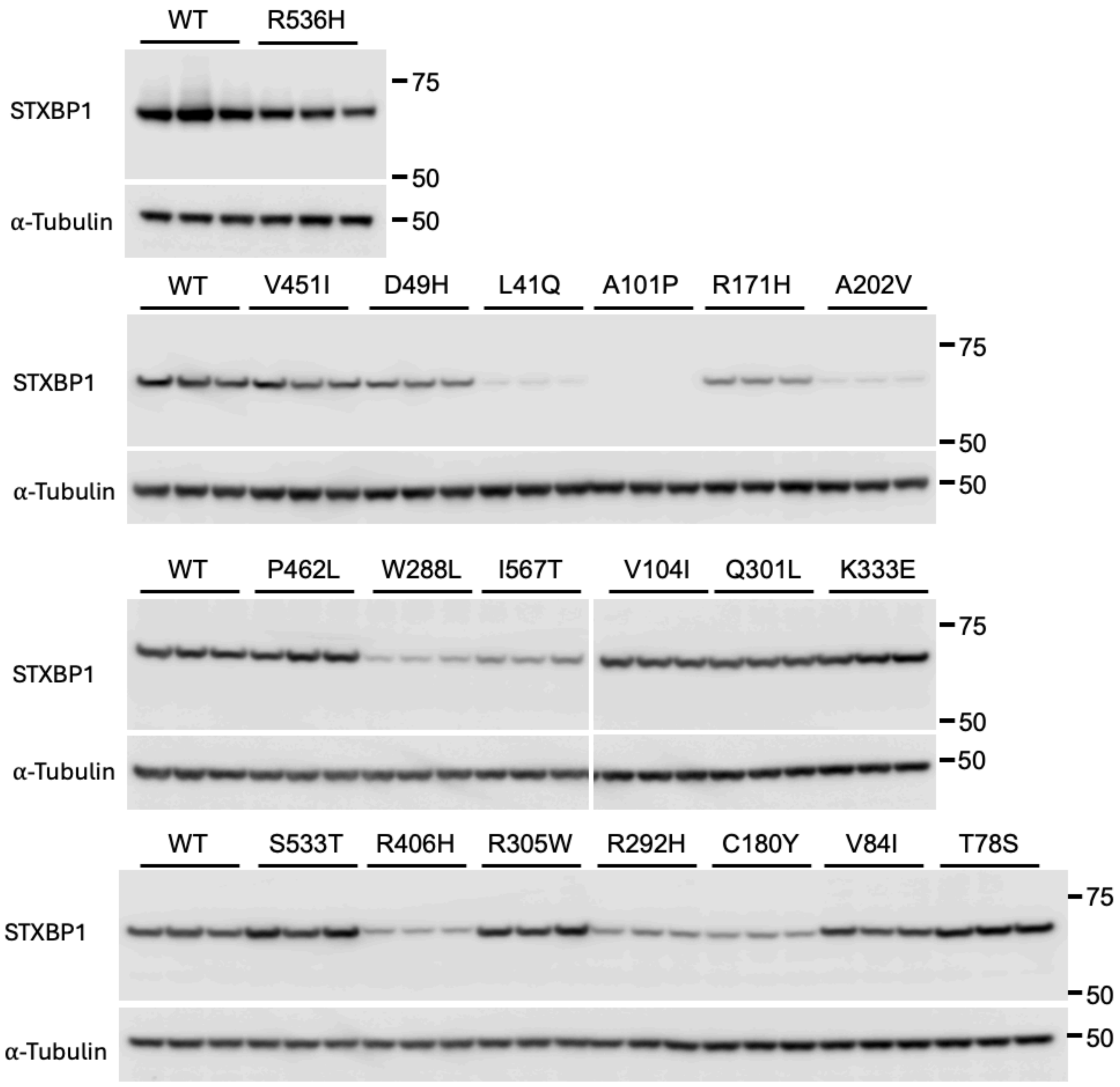

**Supplementary Figure 6: Functional characterization assay for STXBP1: protein abundance assay.**

Cells were transfected with equal amounts of either WT or variant expression plasmids and STXBP1 abundance was measured with an anti-STXBP1 antibody.

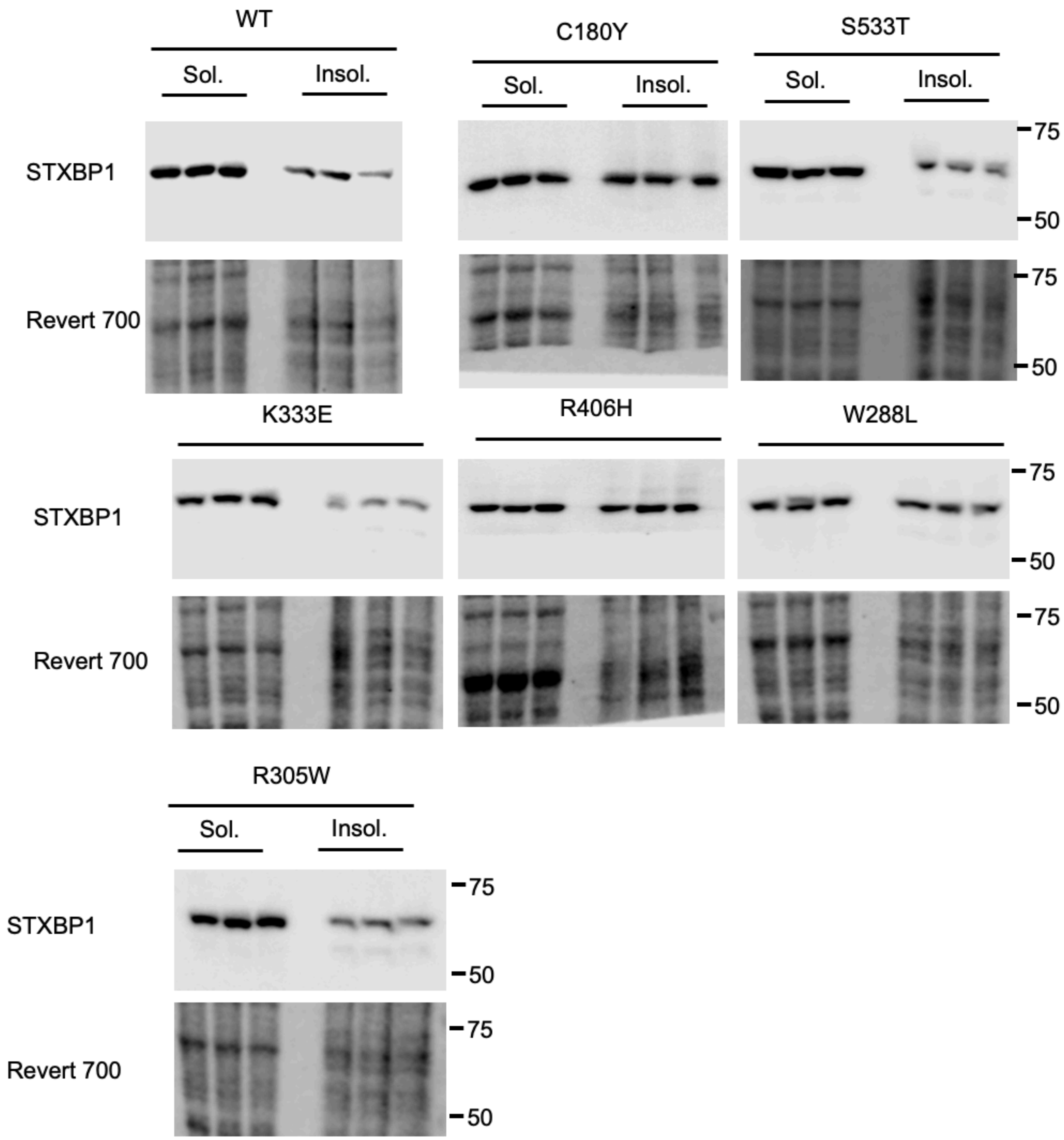
WT
Sol.
Insol.
C180Y
Sol.
Insol.
S533T
Sol.
Insol.
STXBP1
Revert 700
75
50
75
50
K333E
R406H
W288L
STXBP1
Revert 700
75
50
75
50
R305W
Sol.
Insol.
STXBP1
Revert 700
75
50
75
50

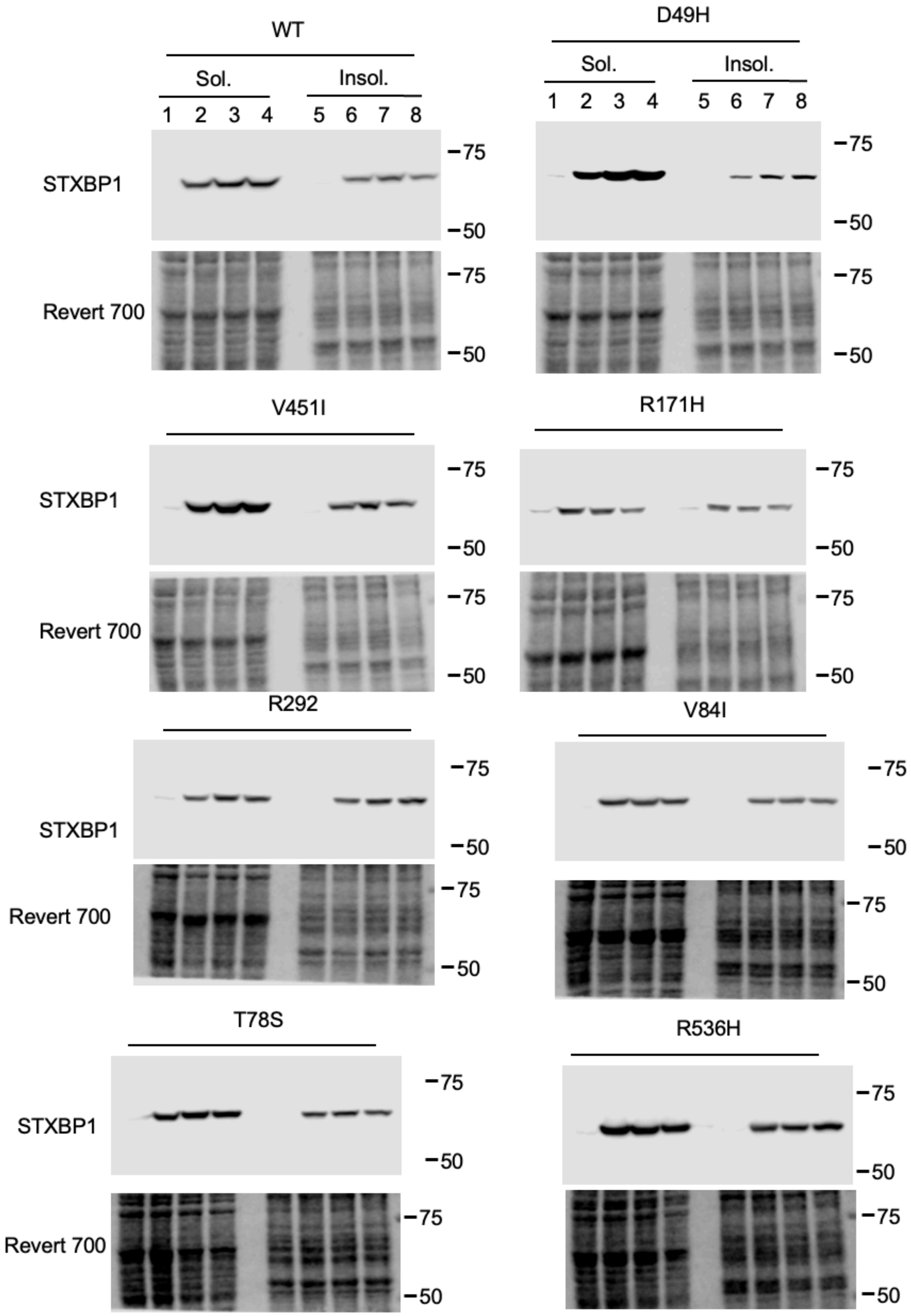
WT
Sol.
Insol.
1 2 3 4 5 6 7 8
STXBP1
Revert 700
75
50
D49H
Sol.
Insol.
1 2 3 4 5 6 7 8
V451I
R171H
R292
V84I
T78S
R536H

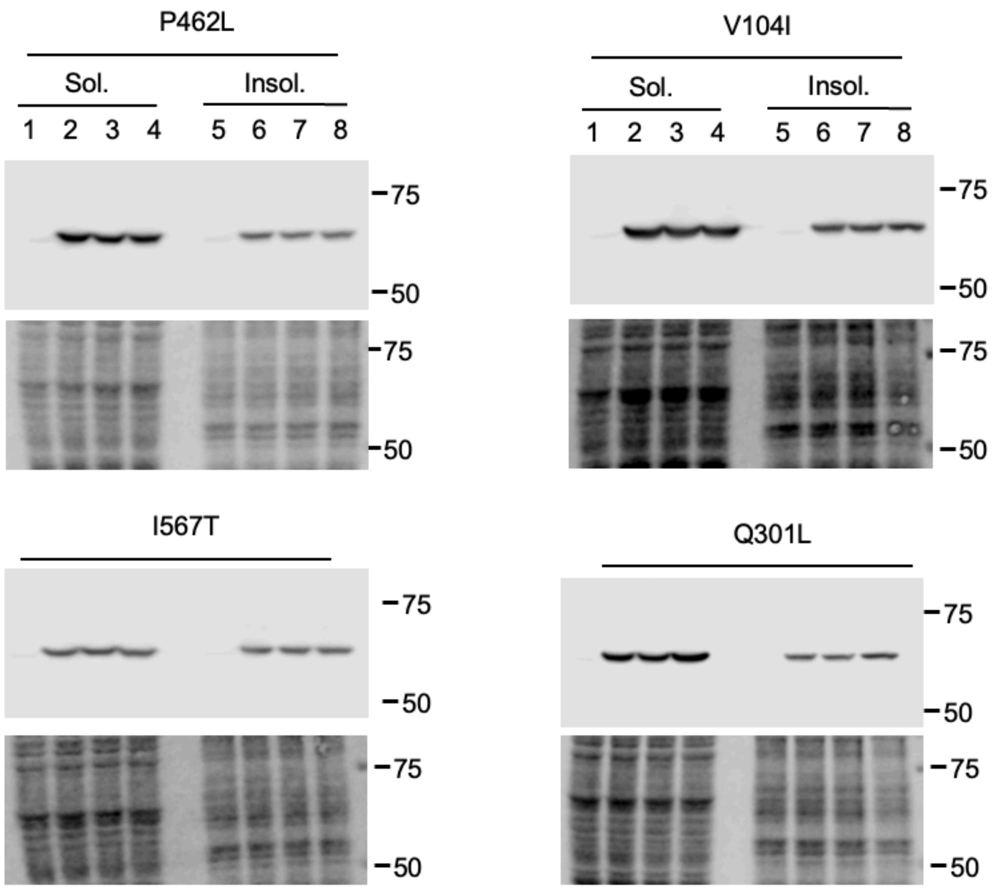

P462L
Sol.
Insol.
1 2 3 4 5 6 7 8
75
50
75
50
V104I
Sol.
Insol.
1 2 3 4 5 6 7 8
75
50
75
50
I567T
75
50
75
50
Q301L
75
50
75
50

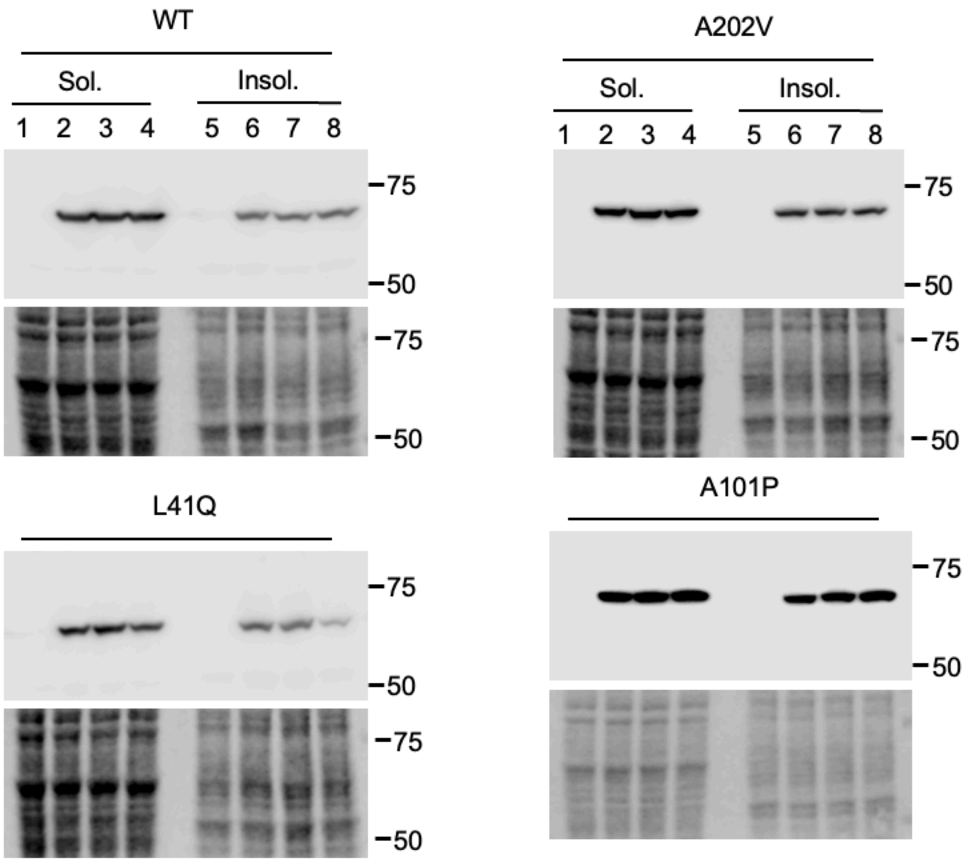

**Supplementary Figure 7: Functional characterization assay for STXBP1: protein solubility assay.**
Cells were transfected with equal amounts of either WT or variant expression plasmids and insoluble and soluble fractions were collected. STXBP1 abundance was measured in each fraction. Revert700 reagent was utilized to confirm loading of protein.

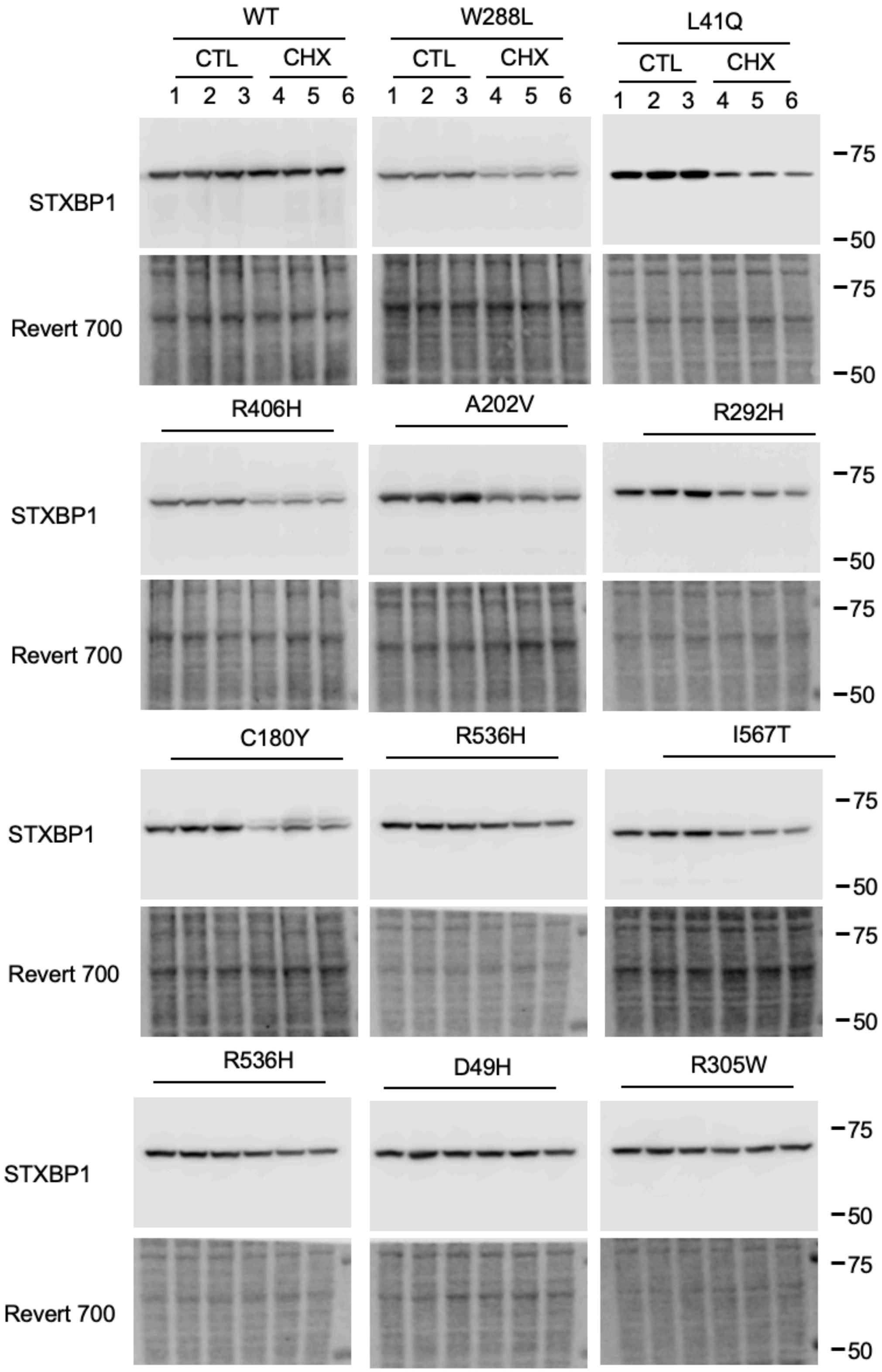

WT
W288L
L41Q
CTL
CHX
1 2 3 4 5 6
STXBP1
Revert 700
75
50
R406H
A202V
R292H
C180Y
R536H
I567T
R536H
D49H
R305W

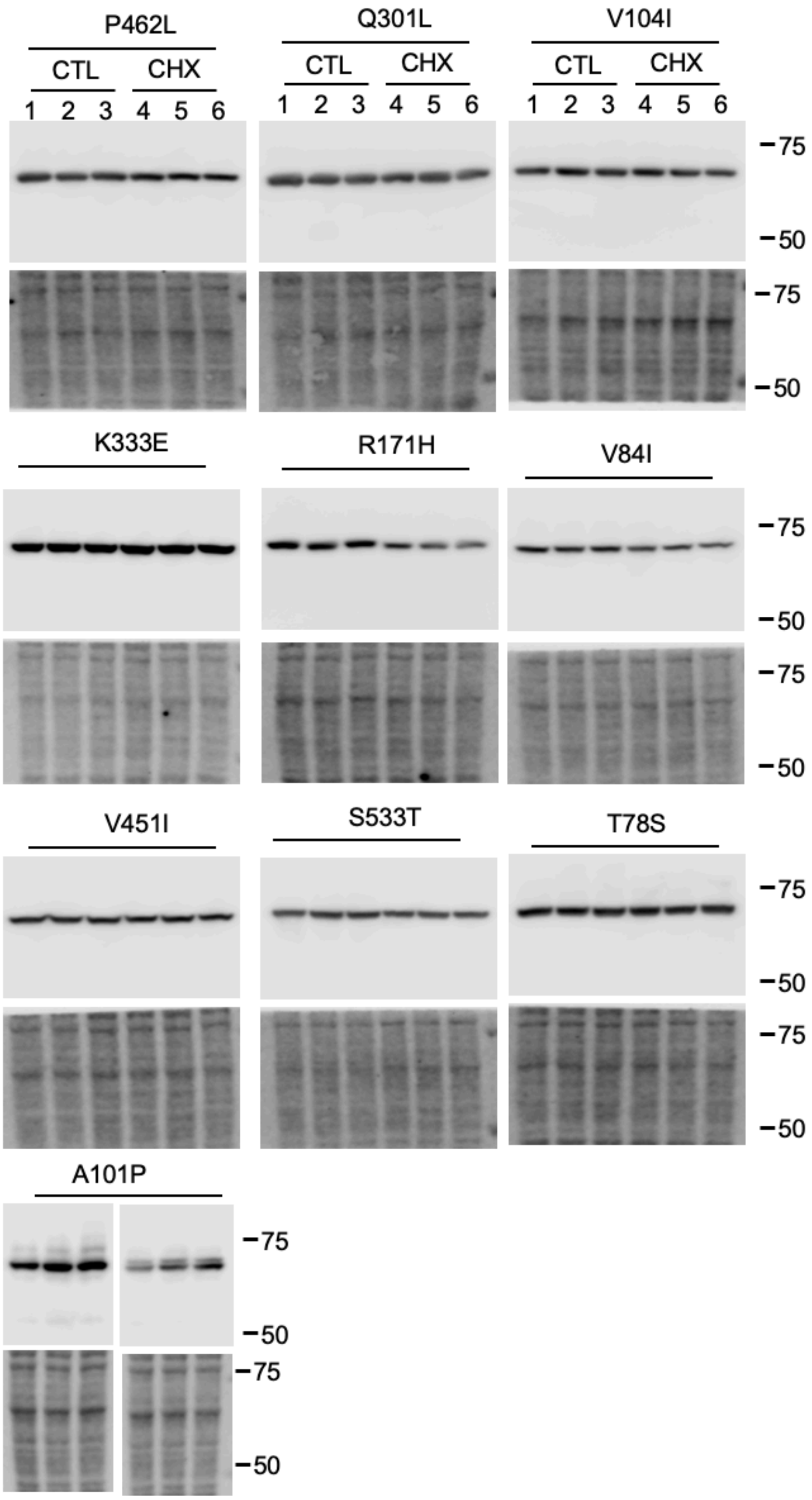

P462L
Q301L
V104I
CTL
CHX
1 2 3 4 5 6
75
50
K333E
R171H
V84I
V451I
S533T
T78S
A101P

**Supplementary Figure 8: Functional characterization assay for STXBP1: protein stability assay.**

Cells were transfected with equal amounts of either WT or variant expression plasmids. Cells were treated with either vehicle (DMSO) or cycloheximide (CHX) for 8 hrs prior to collection of total cell lysate and STXBP1 abundance was measured in each treatment condition. Revert700 reagent was utilized to confirm loading of protein.

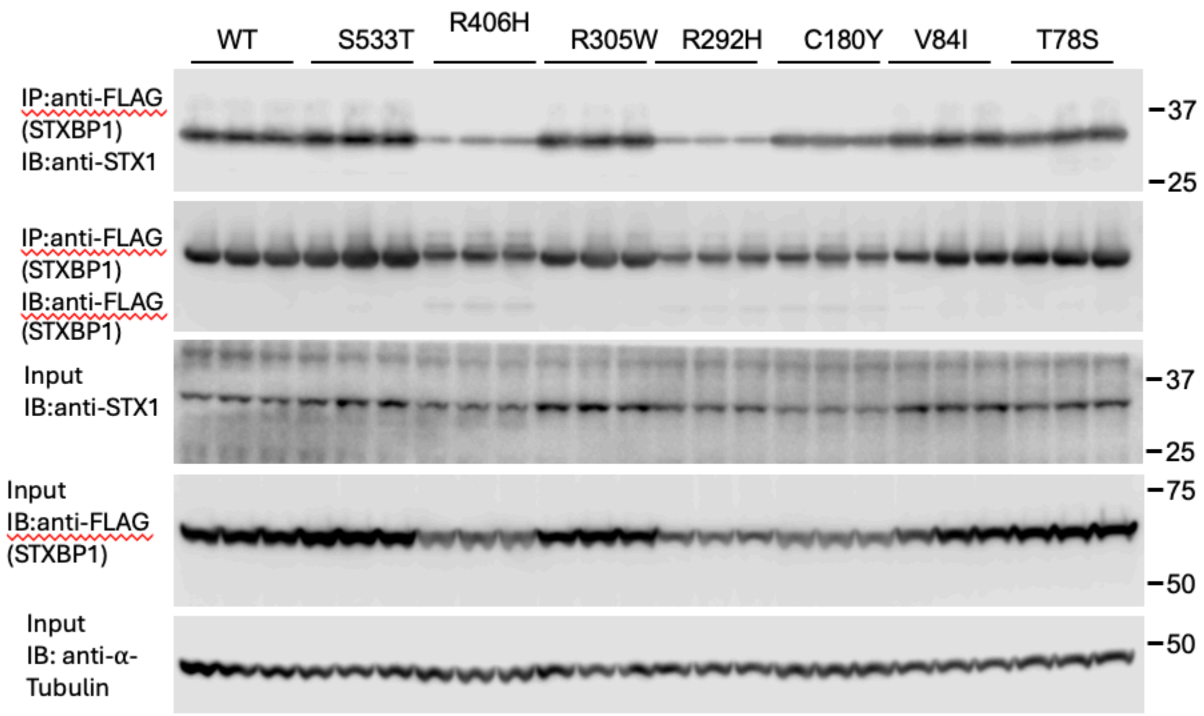

WT
S533T
R406H
R305W
R292H
C180Y
V84I
T78S
IP:anti-FLAG (STXBP1) IB:anti-STX1
IP:anti-FLAG (STXBP1) IB:anti-FLAG (STXBP1)
Input IB:anti-STX1
Input IB:anti-FLAG (STXBP1)
Input IB: anti-α-Tubulin
37
25
37
25
75
50
50

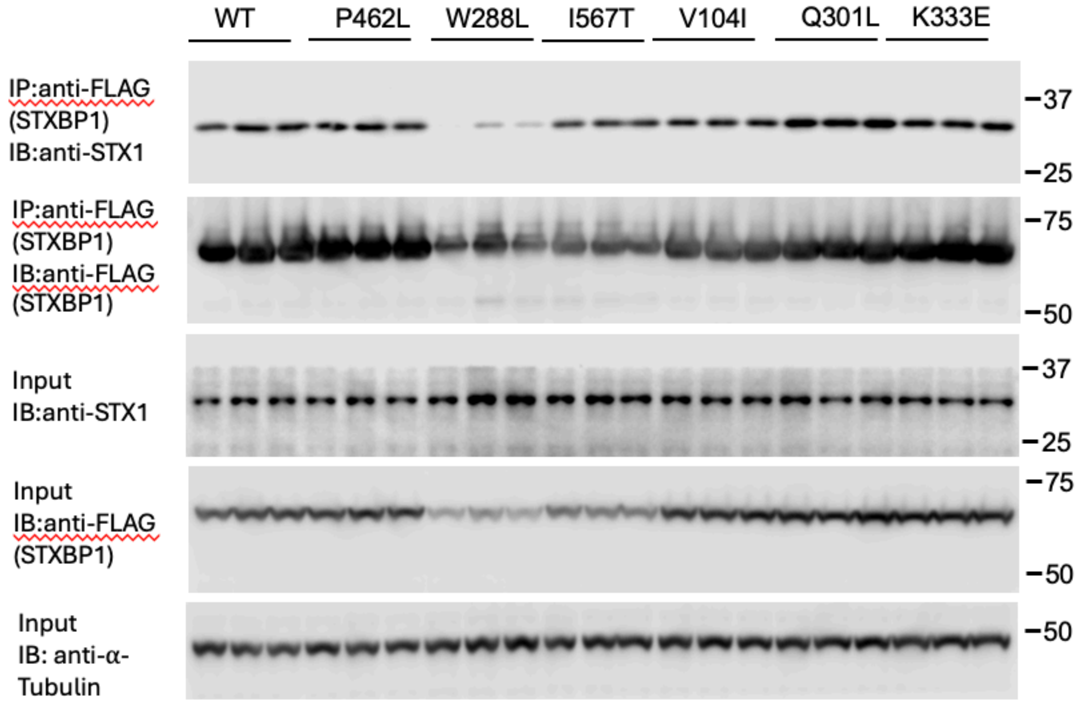

WT
P462L
W288L
I567T
V104I
Q301L
K333E
IP:anti-FLAG (STXBP1) IB:anti-STX1
IP:anti-FLAG (STXBP1) IB:anti-FLAG (STXBP1)
Input IB:anti-STX1
Input IB:anti-FLAG (STXBP1)
Input IB: anti-α-Tubulin
37
25
75
50
37
25
75
50
50

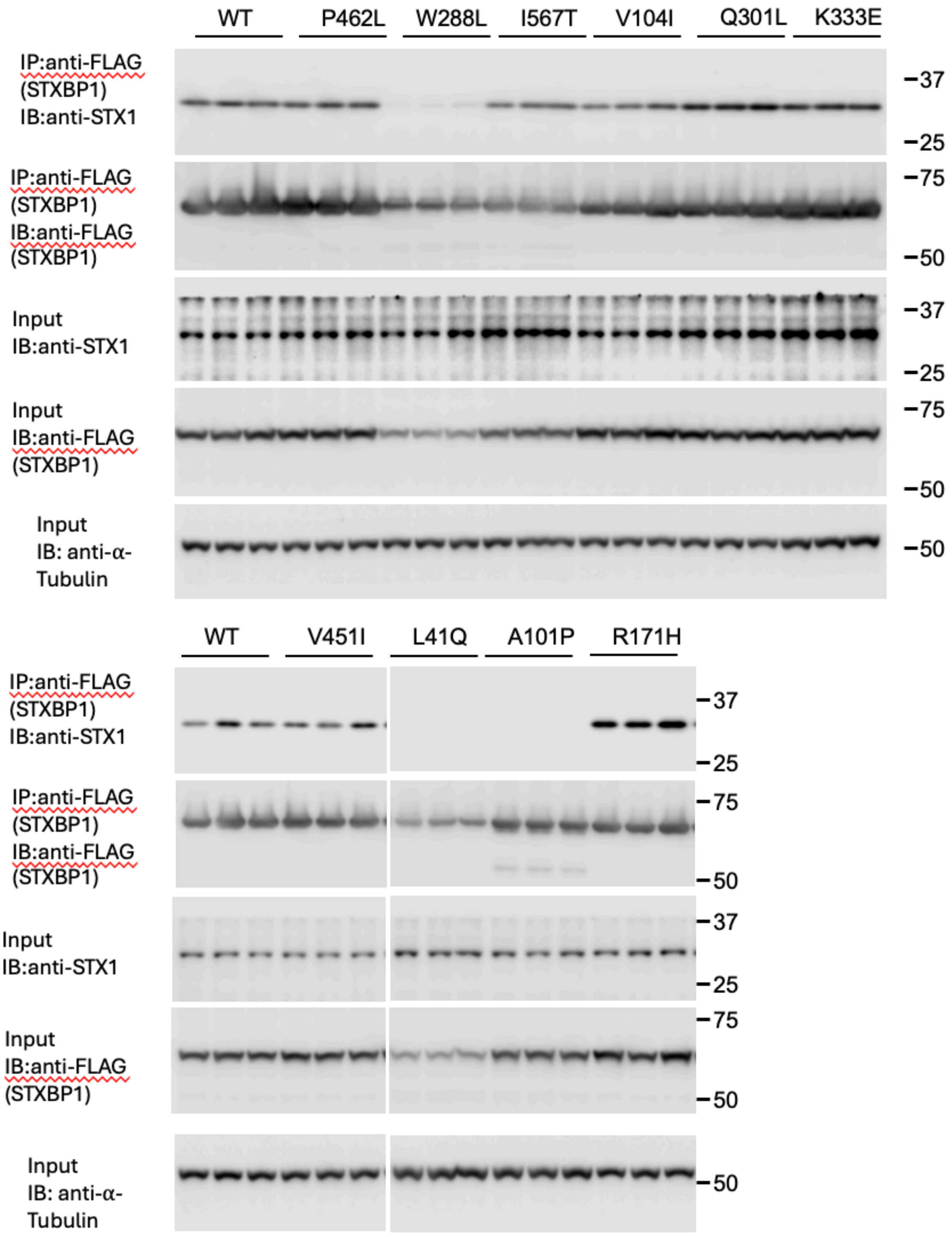
WT
P462L
W288L
I567T
V104I
Q301L
K333E
IP:anti-FLAG (STXBP1) IB:anti-STX1
IP:anti-FLAG (STXBP1) IB:anti-FLAG (STXBP1)
Input IB:anti-STX1
Input IB:anti-FLAG (STXBP1)
Input IB: anti-α-Tubulin
37
25
75
50
WT
V451I
L41Q
A101P
R171H

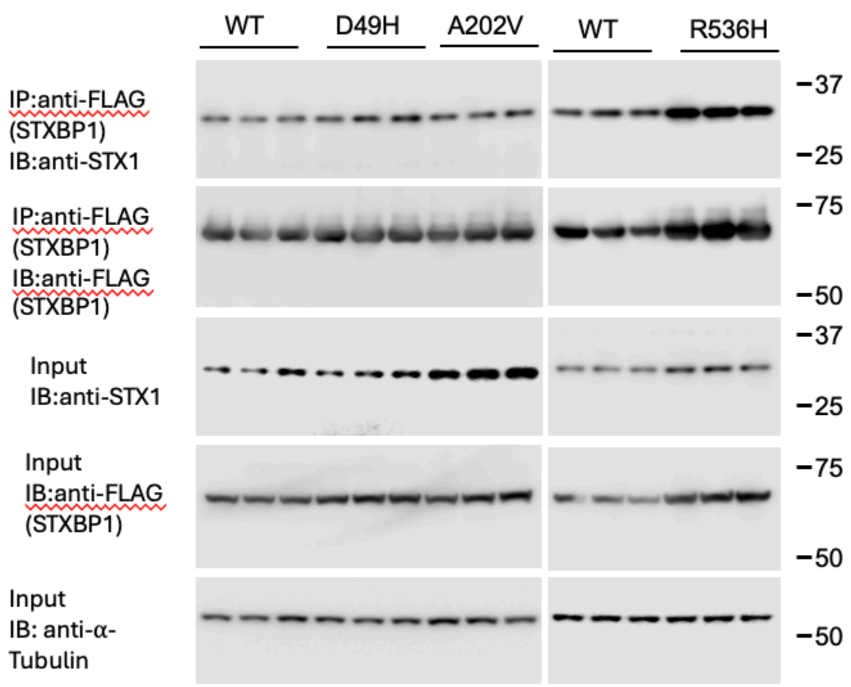

**Supplementary Figure 9: Functional characterization assay for STXBP1 interaction with STX1.**

TRE-T-SNARE cells in 6xwell plates were transfected with different amounts of STXBP1 plasmid DNA to ensure a similar protein expression. Additionally, cells were co-transfected with equal amounts of an STX1 expression plasmid. Interaction was assessed by immunoprecipitation (IP) of STXBP1 followed by immunoblotting (IB) for STX1.

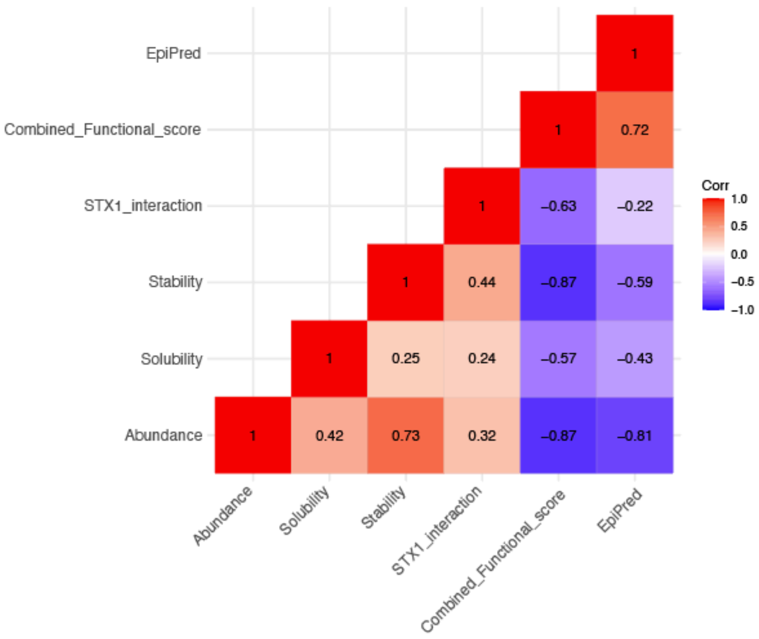

**Supplementary Figure 10: Correlation plot (Pearson) across functional readouts, combined scores and EpiPred**

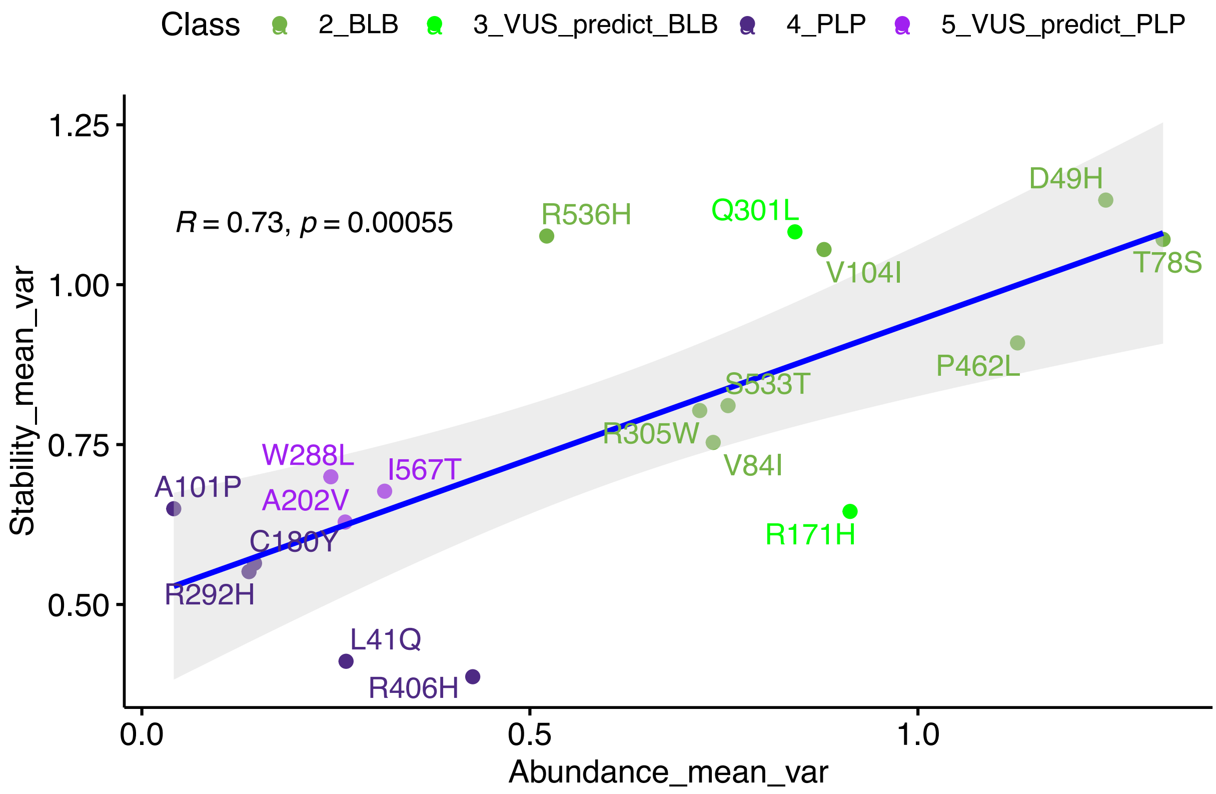
Class
2_BLB
3_VUS_predict_BLB
4_PLP
5_VUS_predict_PLP
R = 0.73, p = 0.00055
Stability_mean_var
Abundance_mean_var
1.25
1.00
0.75
0.50
0.0
0.5
1.0
R536H
Q301L
D49H
V104I
T78S
P462L
S533T
R305W
V84I
R171H
W288L
I567T
A101P
A202V
C180Y
R292H
L41Q
R406H

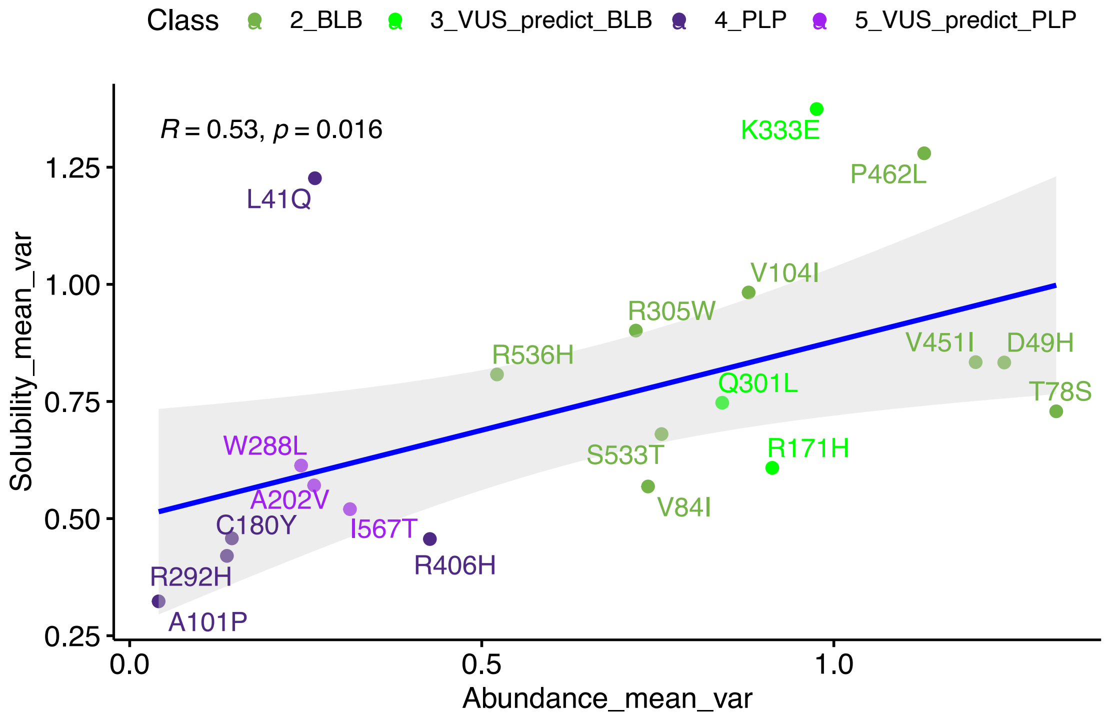
Class
2_BLB
3_VUS_predict_BLB
4_PLP
5_VUS_predict_PLP
R = 0.53, p = 0.016
Solubility_mean_var
Abundance_mean_var
1.25
1.00
0.75
0.50
0.25
0.0
0.5
1.0
K333E
P462L
L41Q
V104I
R305W
R536H
V451I
D49H
Q301L
T78S
S533T
R171H
W288L
A202V
V84I
C180Y
I567T
R406H
R292H
A101P

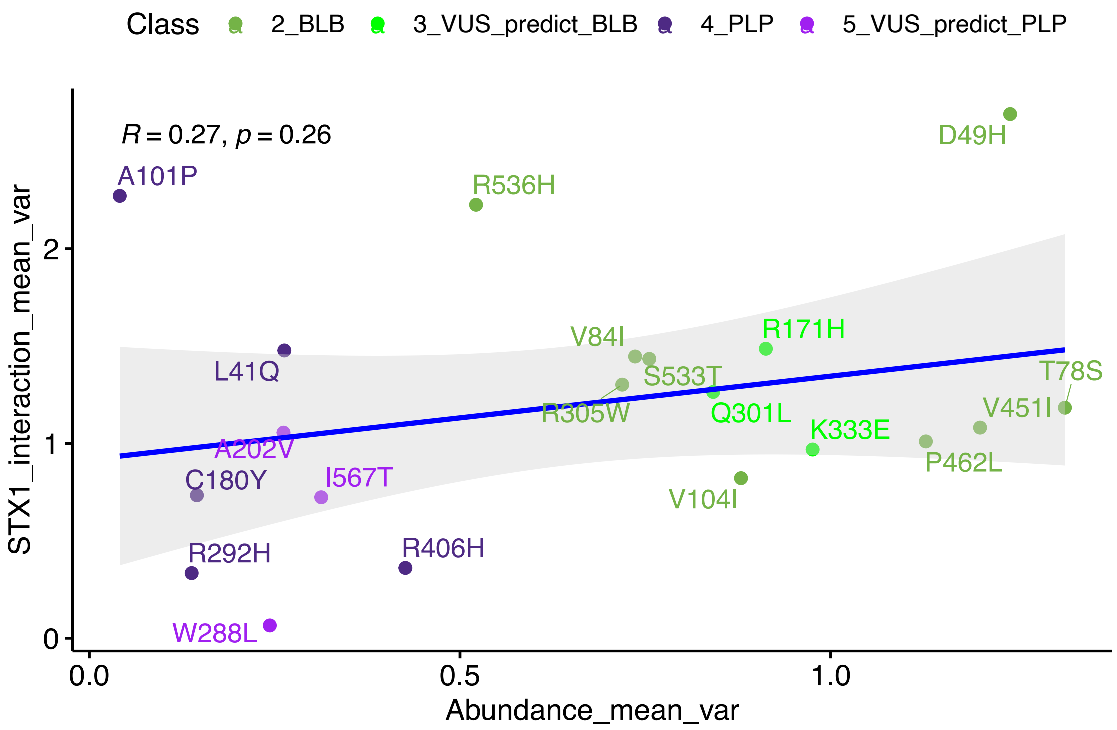

Class
2_BLB
3_VUS_predict_BLB
4_PLP
5_VUS_predict_PLP
R = 0.27, p = 0.26
STX1_interaction_mean_var
Abundance_mean_var
D49H
A101P
R536H
V84I
R171H
L41Q
S533T
T78S
R305W
Q301L
V451I
K333E
A202V
P462L
C180Y
I567T
V104I
R292H
R406H
W288L

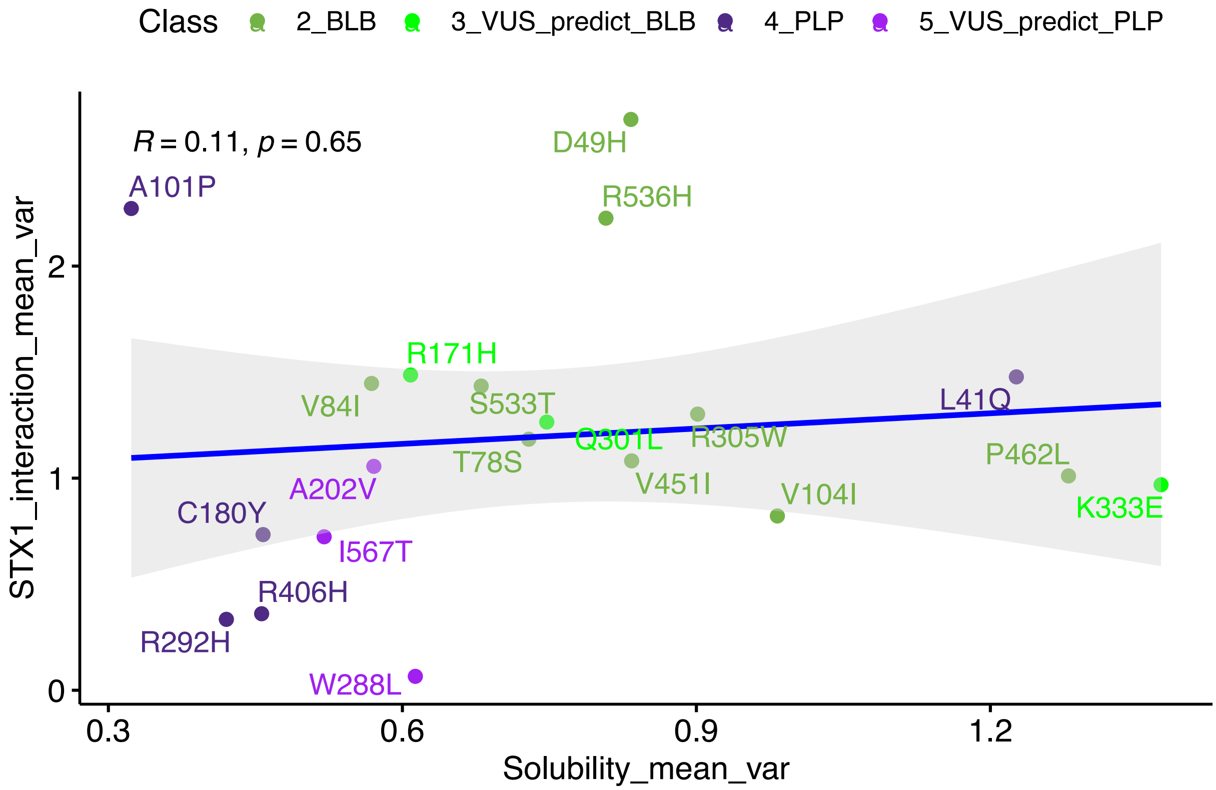

Class
2_BLB
3_VUS_predict_BLB
4_PLP
5_VUS_predict_PLP
R = 0.11, p = 0.65
STX1_interaction_mean_var
Solubility_mean_var
D49H
A101P
R536H
R171H
V84I
S533T
L41Q
Q301L
R305W
T78S
P462L
V451I
V104I
A202V
K333E
C180Y
I567T
R406H
R292H
W288L

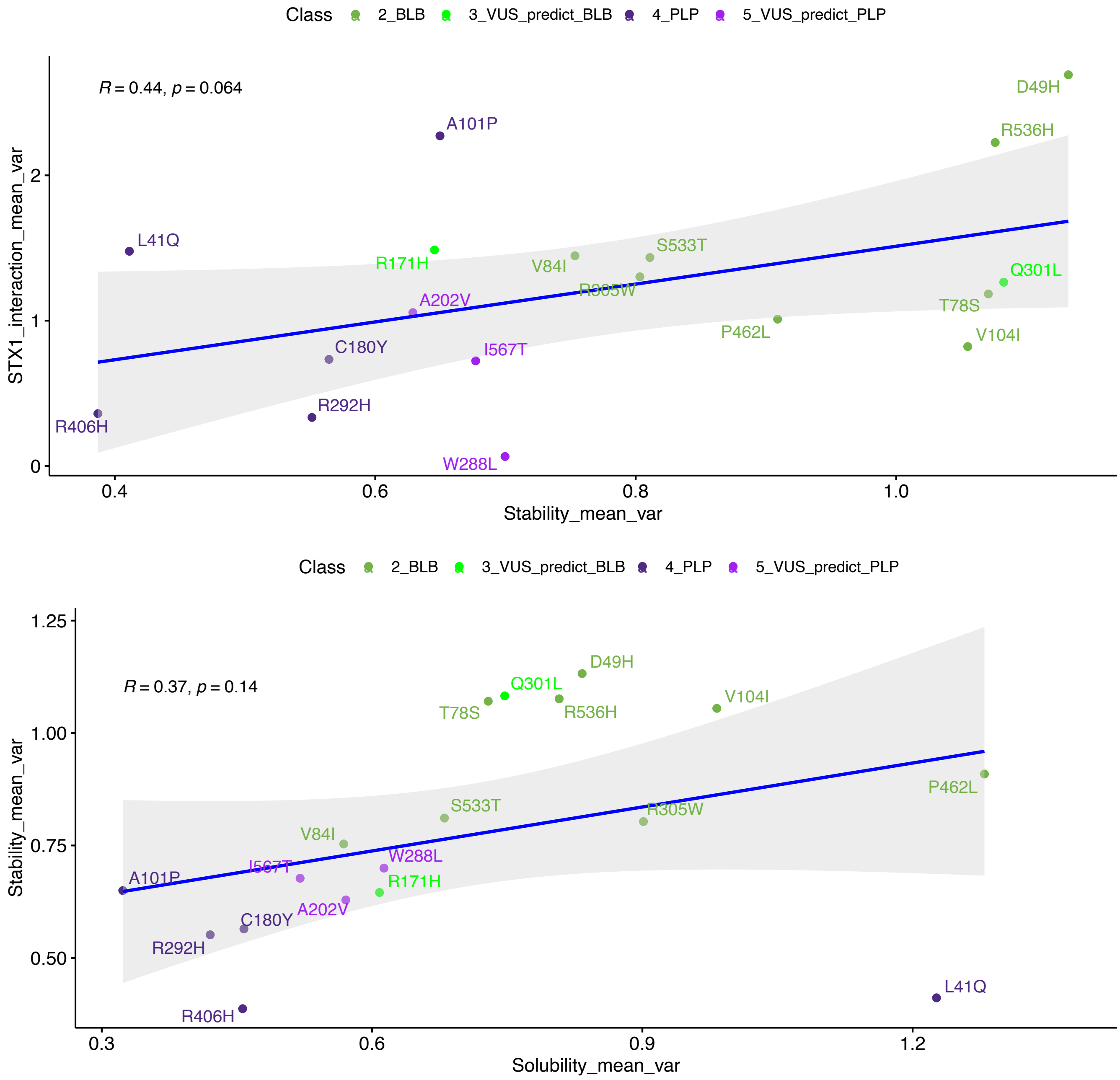

**Supplementary Figure 11: Individual correlation plots (Pearson) across functional readouts**

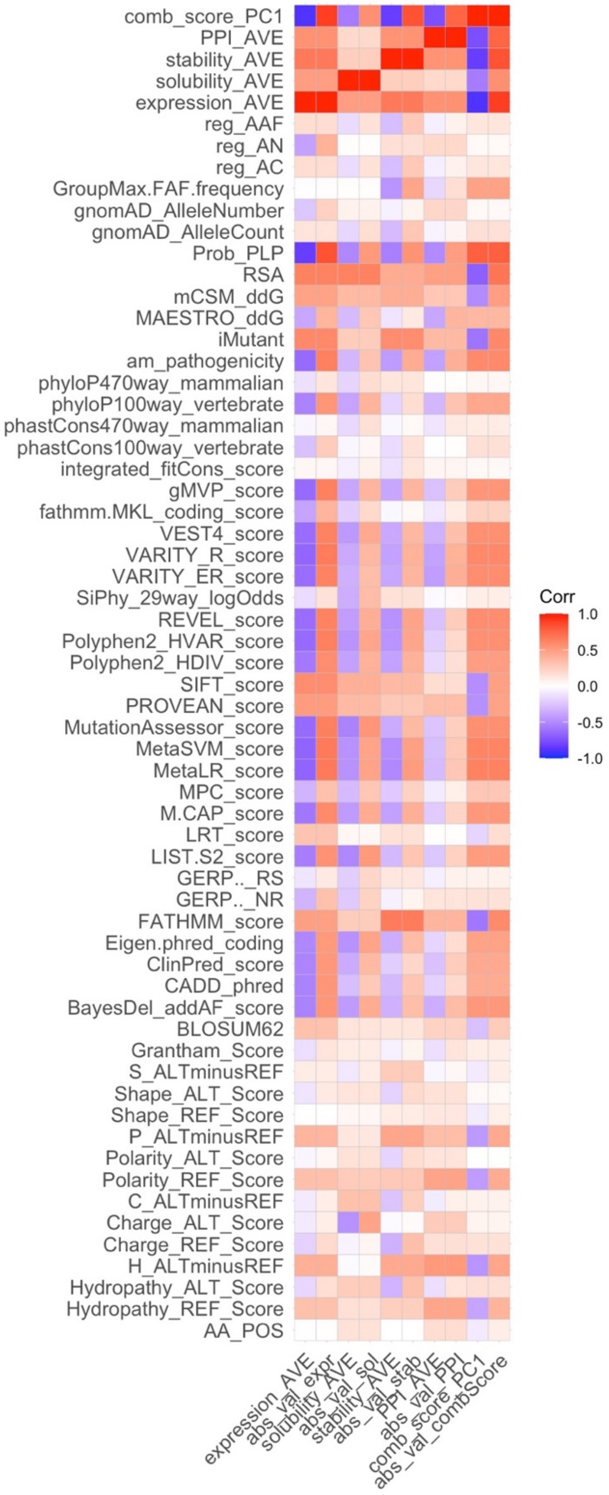

**Supplementary Figure 12: Correlation plot (Pearson) for functional data, EpiPred, and all features used in the EpiPred model as well as common global VEPs**

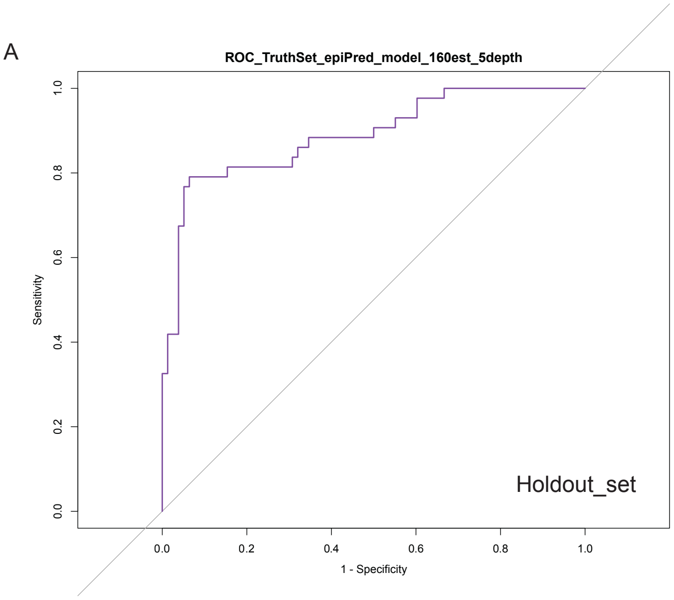

**Supplementary Figure 13: ROC for the EpiPred holdout dataset**

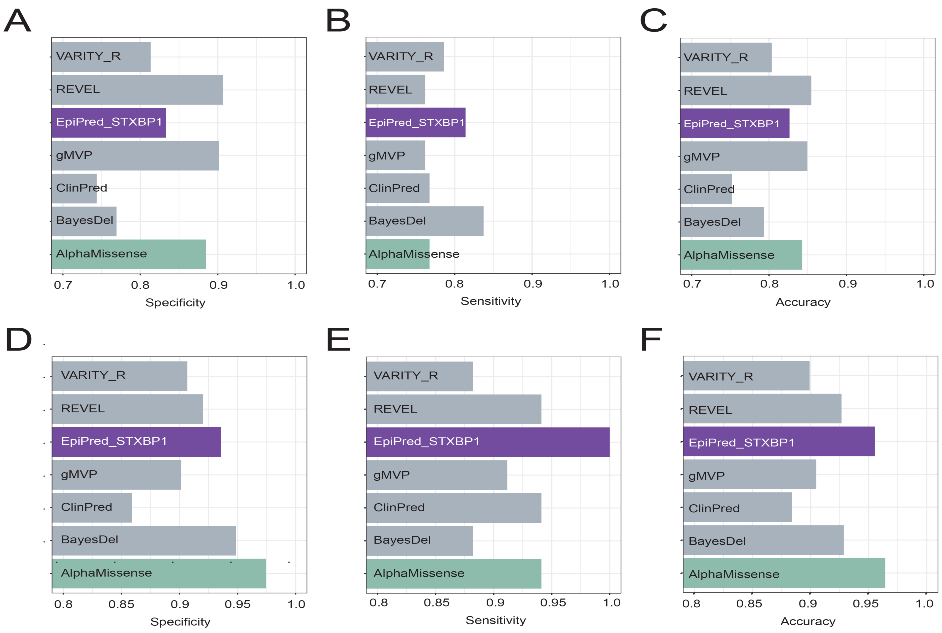

**Supplementary Figure 14: Specificity, sensitivity, and accuracy of EpiPred and other VEPs on the full holdout set with and without the PLP variants (n=9) reclassified as VUS.** (A-C) Metrics on full holdout set. (**D-F**) Metrics on holdout set excluding reclassified variants.

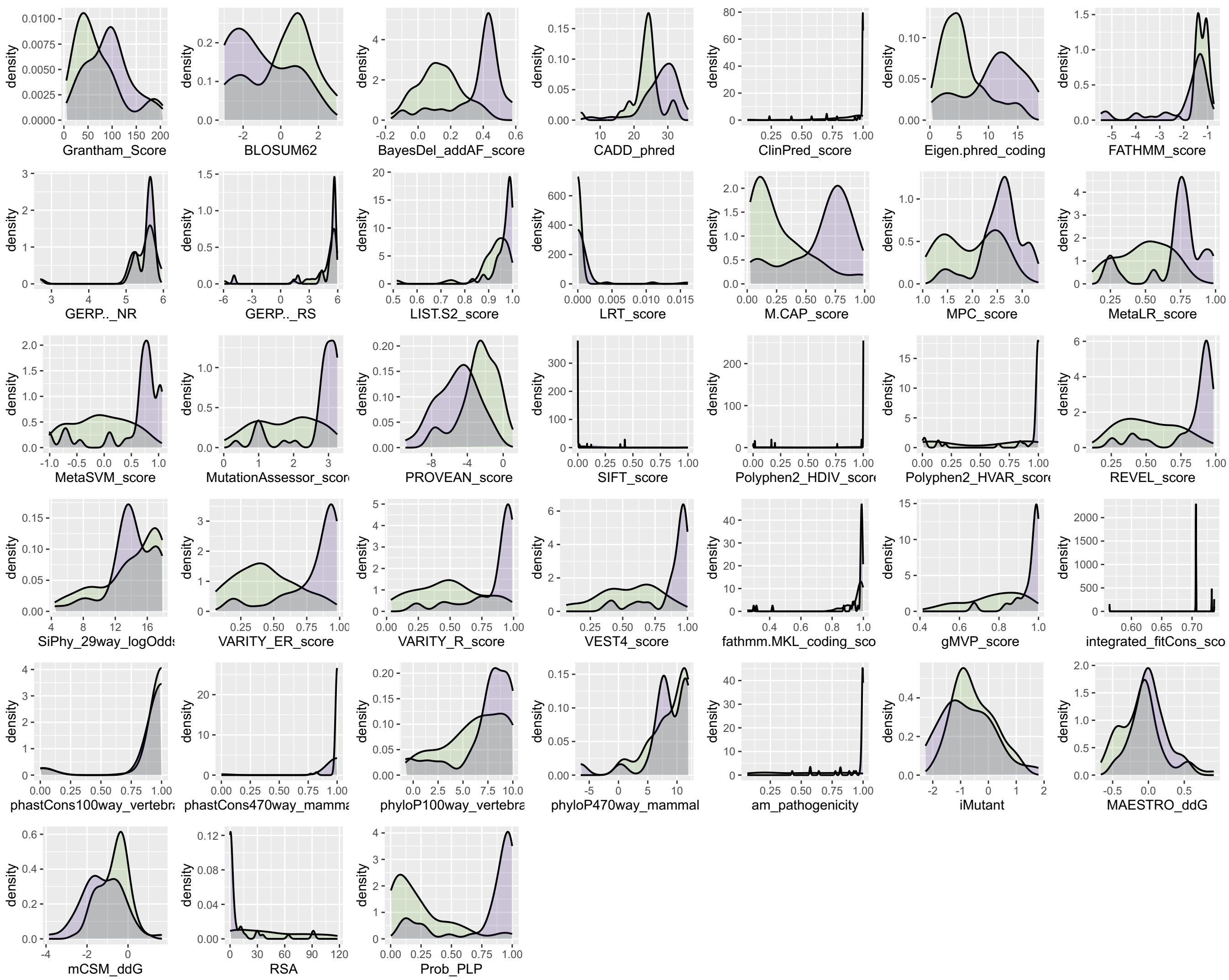

**Supplementary Figure 15: Feature distribution for EpiPred in the holdout set.** Green is class 5 (Holdout set BLB); purple is class 6 (Holdout set PLP). For unabbreviated feature names, please see **Table S3.**

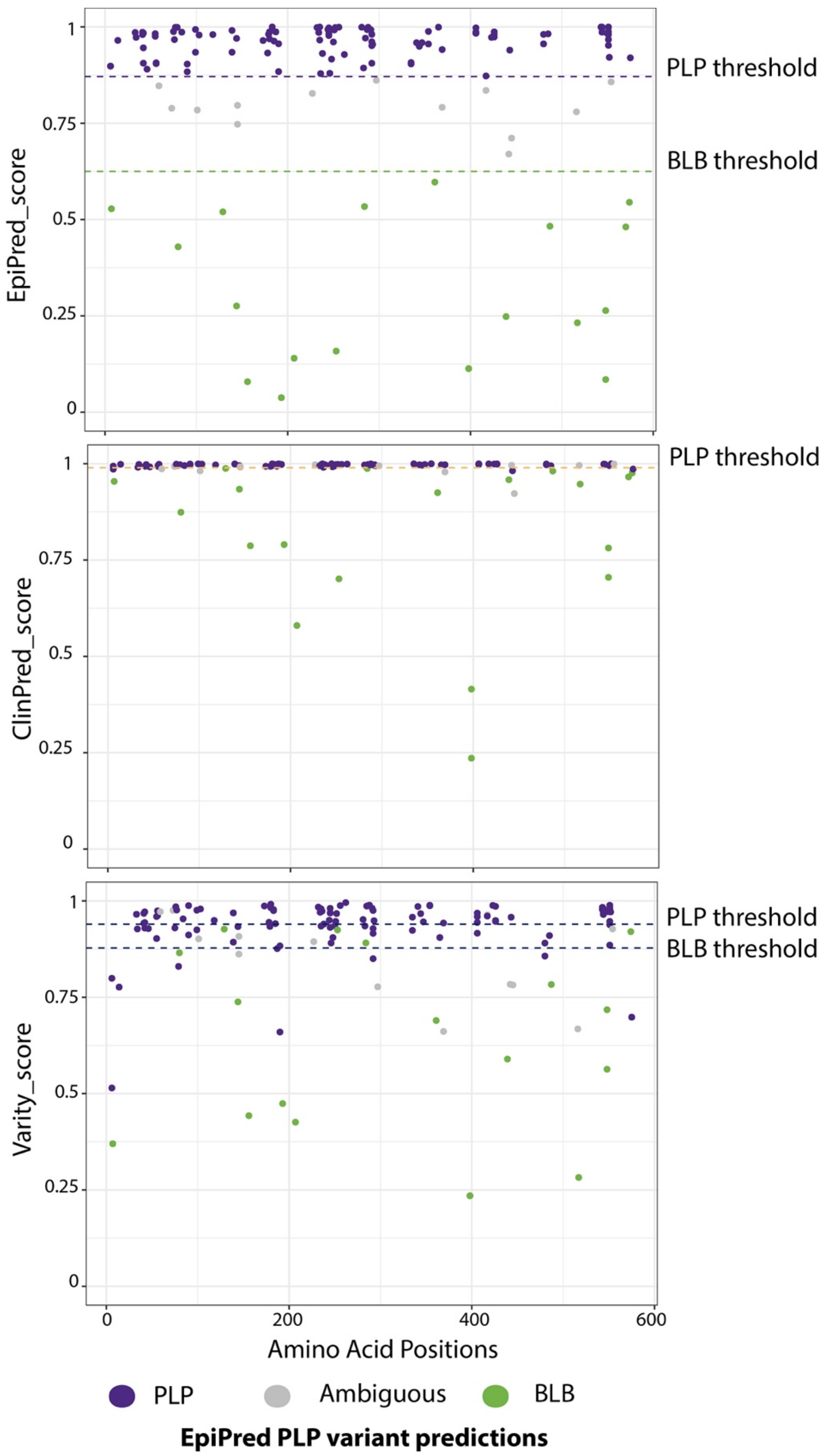

**Supplementary Figure 16: EpiPred and global VEP comparisons**

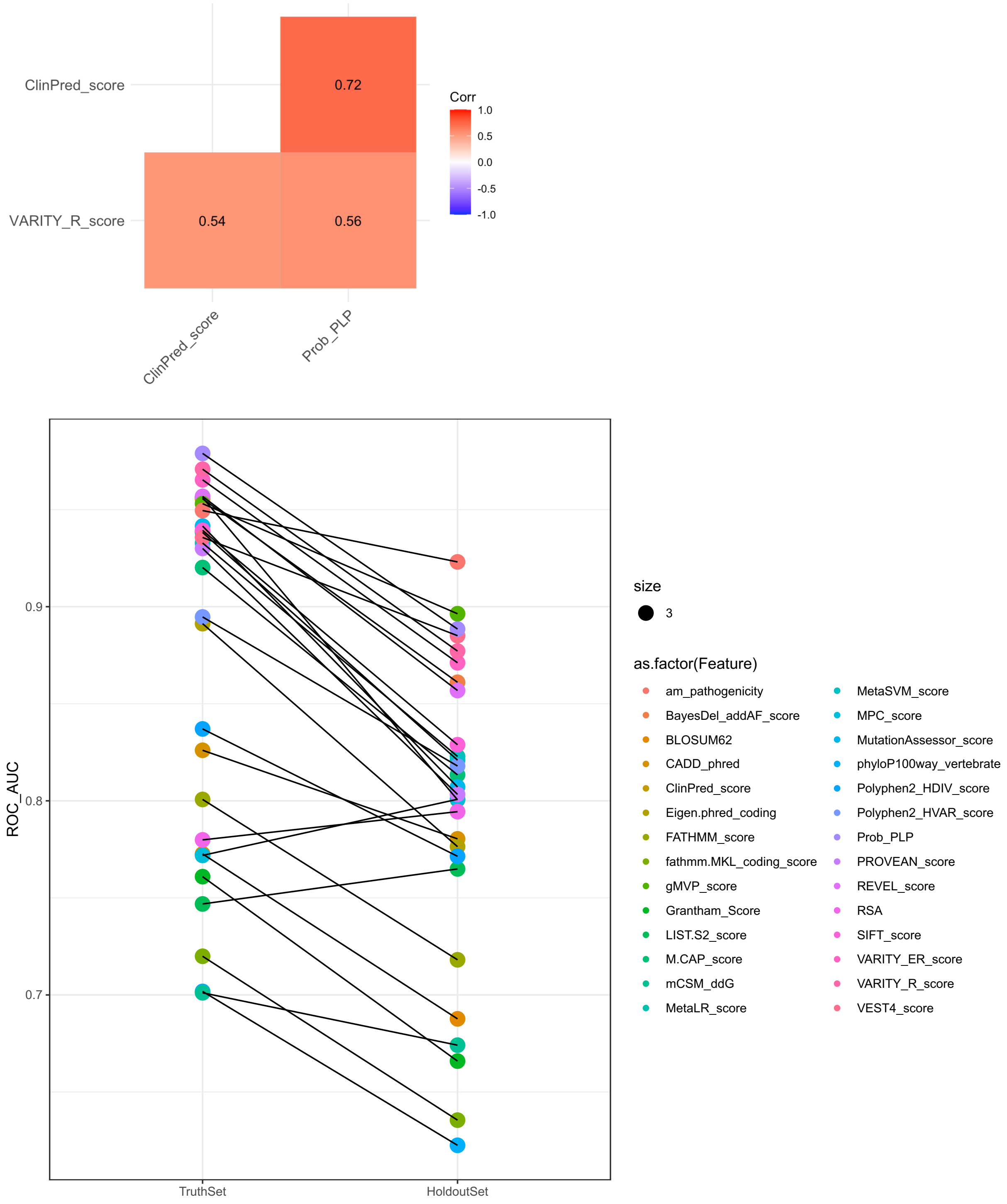

**Supplementary Figure 17: Most VEPs/features have better classification performance on truth set as compared to holdout set.**
ROC$_{AUC}$ shown below, but also true for other model metrics. Filtered for only features/models with at least ROC$_{AUC}$ of 0.7.

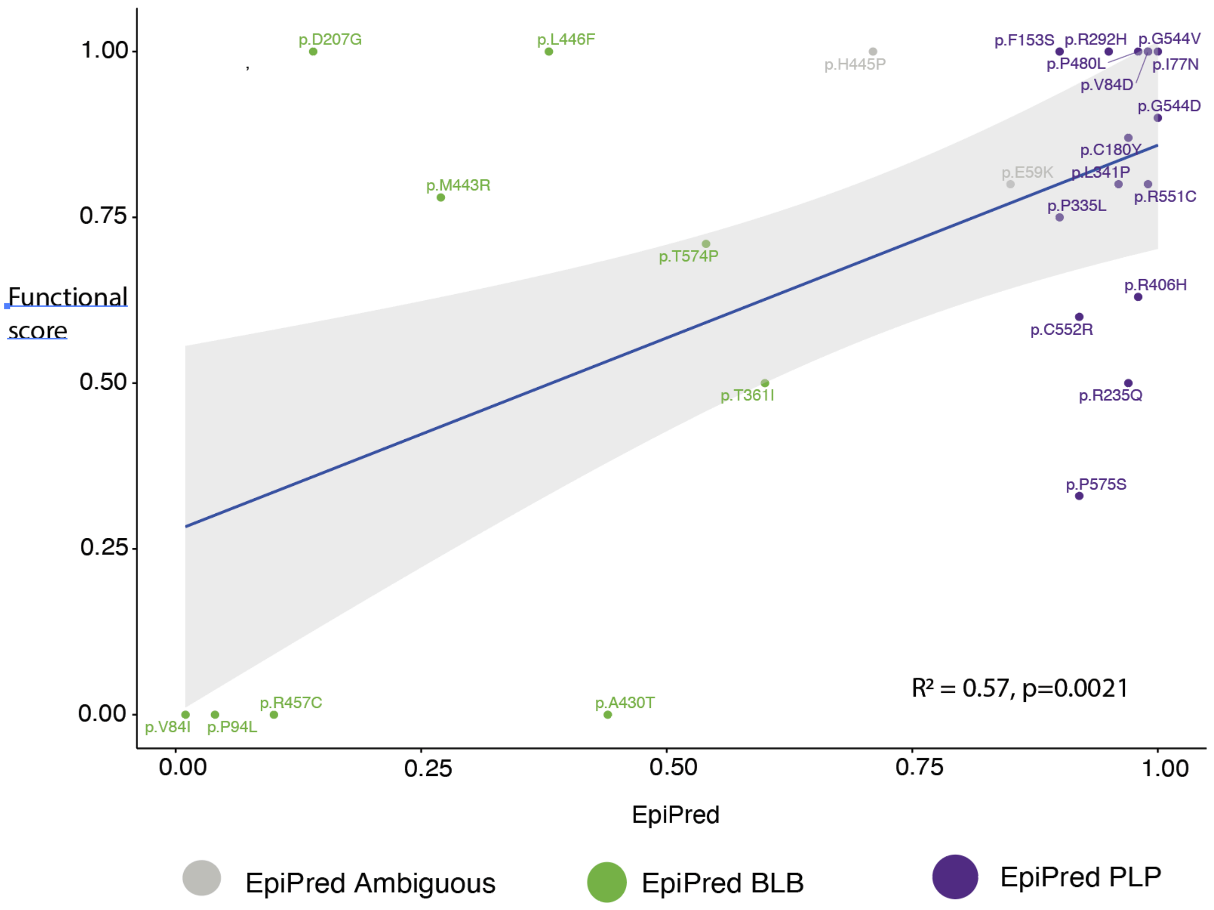

**Supplementary Figure 18: An aggregate functional score from multiple publications, and correlation with EpiPred-PPLP scores**. (r=0.57; p=0.0021 Pearson).

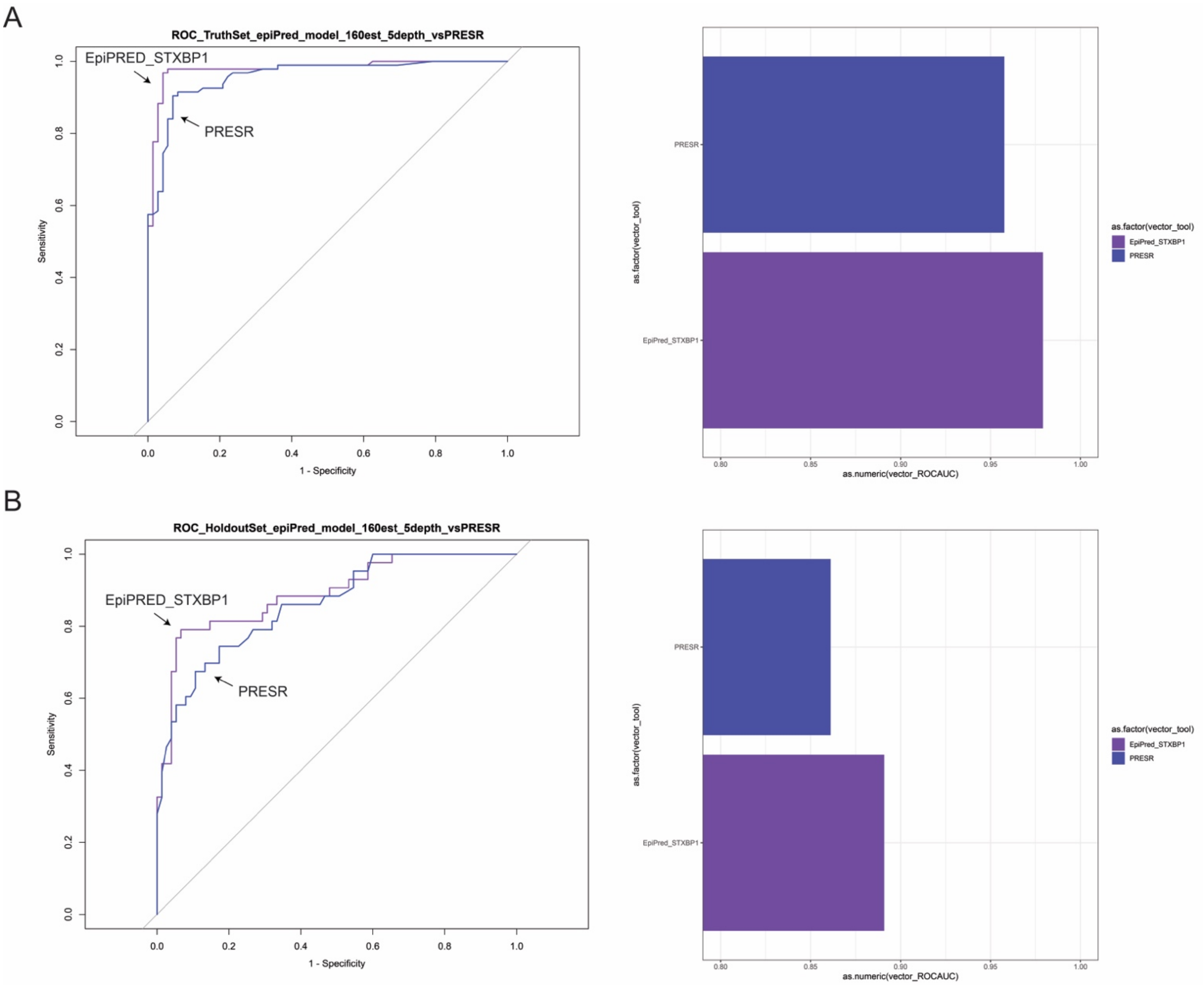

**Supplementary Figure 19: ROC curve and $ROC_{AUC}$ for EpiPRED compared with PRESR.** Plots shown for full truth set (A) and the holdout dataset (B).

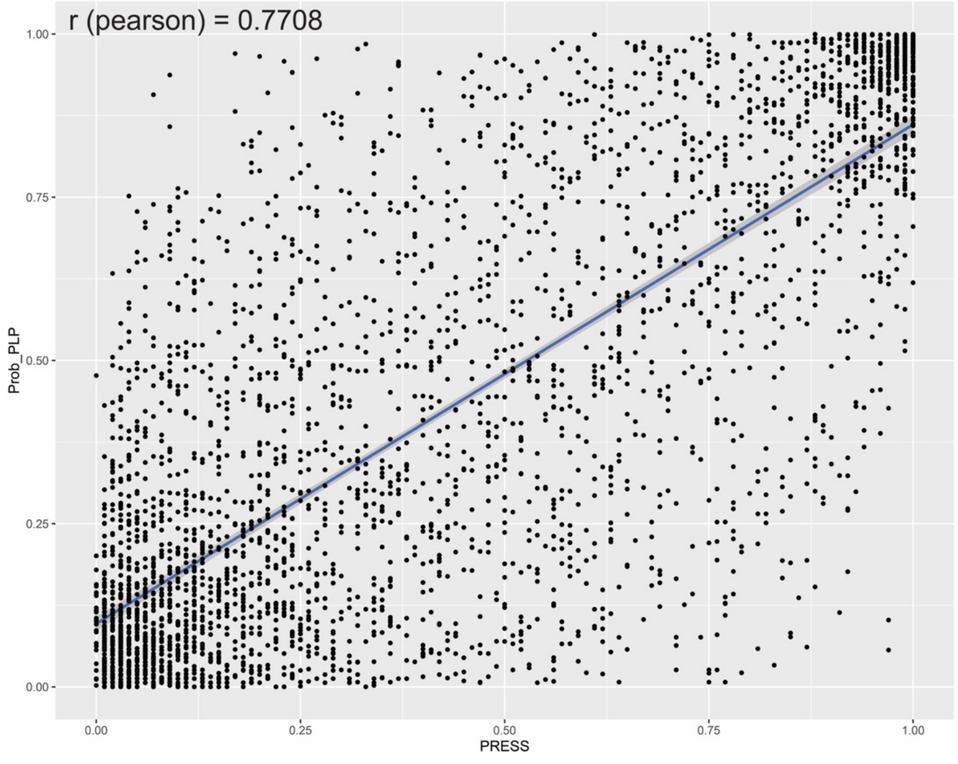

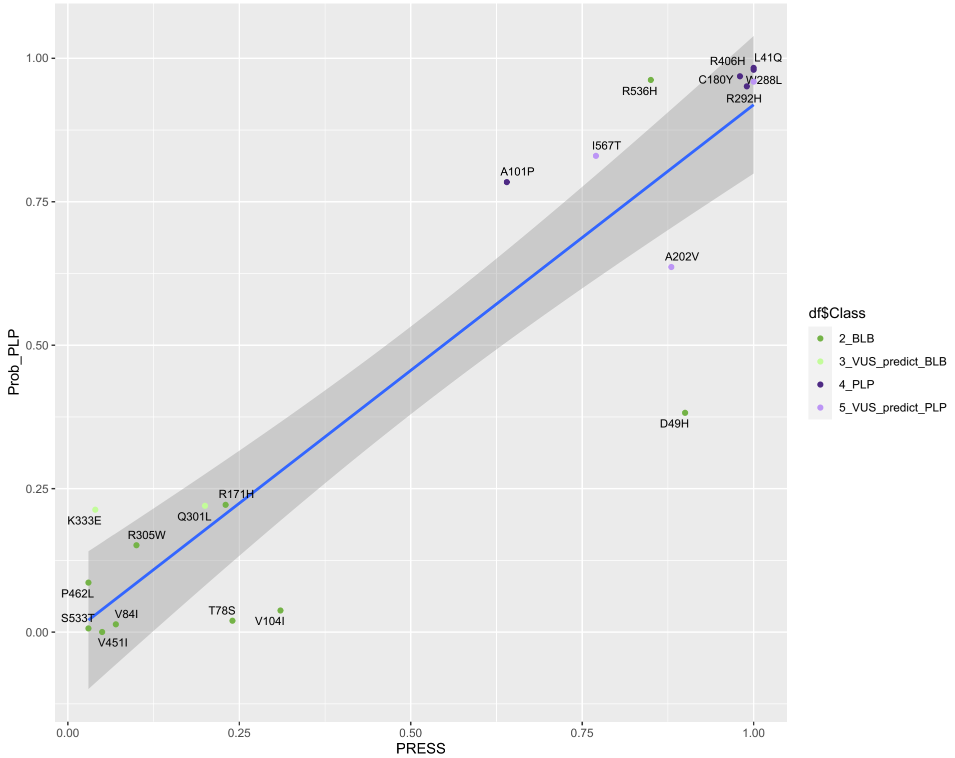

**Supplementary Figure 20: PRESR and EpiPRED are highly correlated.**
(Top) Scatterplot of EpiPred output (Prob_PLP) and PRESR output (PRESS) across full EpiPred dataset including truth set, holdout set, VUS, and simulation variants. (Bottom) Scatterplot for a subset of variants with functional data (n=20) shown.
